# Structural dynamics of the methyl-coenzyme M reductase active site are influenced by coenzyme F_430_ modifications

**DOI:** 10.1101/2024.01.07.574536

**Authors:** Marcelo D. Polêto, Kylie D. Allen, Justin A. Lemkul

## Abstract

Methyl-coenzyme M reductase (MCR) is a central player in methane biogeochemistry, governing methanogenesis and the anaerobic oxidation of methane (AOM) in methanogens and anaerobic methanotrophs (ANME), respectively. The prosthetic group of MCR is coenzyme F_430_, a nickel-containing tetrapyrrole derivative. Additionally, a few modified versions of F_430_ have been discovered, including the 17^2^-methylthio-F_430_ (mt-F_430_) that functions with ANME-1 MCR. This study employs molecular dynamics (MD) simulations to unravel the intricacies of the active-site dynamics of MCR from *Methanosarcina acetivorans* and ANME-1 when bound to the canonical F_430_ compared to 17^2^-thioether coenzyme F_430_ variants and substrates for methane formation. Overall, our simulations indicate that each MCR active site is optimized for a given version of F_430_ and support the importance of the Gln to Val substitution in accommodating the 17^2^ methylthio modification. Notably, modifications in the 17^2^ position disrupt the canonical coordination among cofactors in *M. acetivorans* MCR, implicating structural perturbations, but evidence of active site reorganization to maintain substrate positions suggest that the modified F_430_s could be accommodated in a methanogenic MCR. We additionally report the first quantitative estimate of MCR intrinsic electric fields pivotal in driving methane formation. Our results suggest that the electric field aligned along the CH_3_-S-CoM thioether bond facilitates homolytic bond cleavage, coinciding with the proposed catalytic mechanism. Structural perturbations, however, weaken and misalign these electric fields, emphasizing the significance of the active site structure in maintaining their integrity. In conclusion, our results deepen the understanding of MCR active-site dynamics, the enzyme’s organizational role in intrinsic electric fields for catalysis, and the interplay between active site structure and electrostatics. This work not only advances our comprehension of MCR functionality but also provides a foundation for future investigations employing sophisticated models to capture the complex electronic properties of MCR active sites quantitatively.

## Introduction

Methanogens are a diverse group of archaea found in a wide range of anaerobic environments including marine and freshwater ecosystems, anaerobic digesters, and animal microbiomes ^1–3^. These organisms possess a unique energy metabolism known as methanogenesis, which is responsible for at least 70% of global methane emissions. The methane-forming step of methanogenesis is catalyzed by methyl-coenzyme M reductase (MCR), in which methyl-coenzyme M (CH_3_-S-CoM) and coenzyme B (HS-CoB) are converted to methane and the CoM-S-S-CoB heterodisulfide (**Figure 1A**)^4^. MCR is a dimer of heterotrimers with a α_2_β_2_γ_2_ configuration, and two active sites that each harbor the nickel-hydrocorphin prosthetic group, coenzyme F_430_ ^5^ (**Figure 1A**). The current working mechanism^6^ involves radical chemistry, which is supported by several computational studies^7–10^ (**Figure 1B**). Ni(I) induces homolytic cleavage of the methyl-sulfur bond of CH_3_-S-CoM to generate a transient methyl radical and a bond between Ni(II) and S-CoM. The methyl radical abstracts the hydrogen atom from HS-CoB to produce methane and •S-CoB, which reacts with the Ni-bound CoM to generate a disulfide radical anion. One-electron transfer to the Ni then releases the heterodisulfide and regenerates Ni(I)^6^. Recently, an alternate binding mode for CH_3_-S-CoM was proposed in which the sulfonate – instead of the thioether – is coordinated to the Ni of F_430_^11^. Thus, although the radical mechanism is conserved overall, this orientation would require two steps involving long-range electron transfer through CoM.

**Figure 1.**
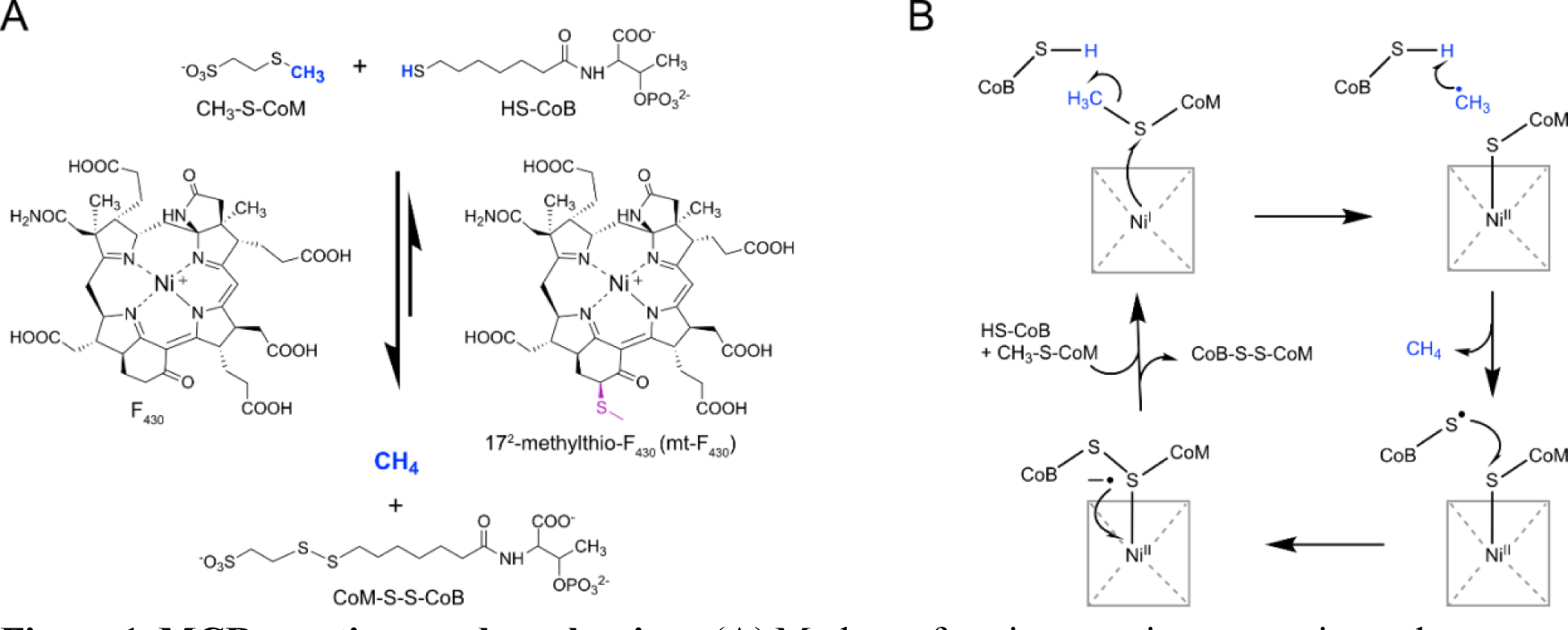
MCR reactions and mechanism. (A) Methane-forming reaction occurs in methanogens and the reverse methane-oxidation reaction occurs in ANME. The structure of F_430_ is shown as well as the 17^2^-methylthio-F_430_ found in ANME-1. (B) Working mechanism involving one-electron chemistry with a methyl radical intermediate (Wongnate et al, 2016).

Functioning in the reverse direction, MCR also catalyzes the first step in the anaerobic oxidation of methane (AOM) carried out by archaea related to methanogens known as anaerobic methanotrophs (ANME)^12,13^. Interestingly, one group of ANME organisms – *Ca*. Methanophagales (ANME-1) – contain a modified form of F_430_ (**Figure 1C**). The structure was assigned as 17^2^-methylthio-F ^14^ (hereafter referred to as mt-F_430_), which was later confirmed in the active site of an ANME-1 MCR crystal structure^15^. The F_430_ modification is accommodated by a valine residue in place of the corresponding glutamine residue (which is often methylated at the 2-position) found in most other MCRs. In addition to mt-F_430_, a handful of modified F_430_s were discovered in small molecule extracts from methanogens^16^ and a crystal structure of the MCR homolog in ethane-oxidizing archaea revealed dimethyl-F_430_ (methyl groups at the 17 and 17^2^ positions)^17^. The roles of F_430_ modifications with respect to MCR structure and catalysis remain unclear. Recently, computational studies revealed that F_430_, compared to its biosynthetic precursors, exhibits distinct chemical properties and, thus, reactivity in the methane-forming reaction^18^. Notably, the reduction potential becomes progressively and substantially higher going from early biosynthetic precursors to the final F_430_, leading to the conclusion that modifications to the macrocyclic ring system have a large impact on the redox properties of the Ni. Furthermore, the Ni(II)-S bond of F_430_-Ni(II)-S-CoM was demonstrated to be strongest for F_430_ compared to its precursors, which stabilizes the key transition state promoting CH_3_-S bond cleavage (**Figure 1B**). Thus, additional modifications to F_430_ are expected to influence the reactivity of the coenzyme.

Here, we report the first molecular dynamics (MD) simulations of MCR. We investigated the MCR from a model methanogen, *Methanosarcina acetivorans*, as well as an ANME-1 MCR, in the presence of the canonical F_430_ and two modified versions of F_430_. Our goal was to describe the active site conformational dynamics and how subtle differences amongst the MCR of each organism could contribute to catalysis as well as preferences for specific versions of F_430_. We also investigated the intrinsic electric field driving catalysis in MCR systems to better understand if modification of F_430_ may impact catalysis. Our approach provides new insights into the conformational dynamics of MCR active sites, the relationship between modified F_430_ cofactors and coordination among cofactors, and how MCR is able to organize intrinsic electric fields to facilitate the catalysis of methane formation.

## Methods

### Parametrization of cofactors

Parameters for methyl coenzyme M (CH_3_-S-CoM) and coenzyme B (HS-CoB) were obtained from the CHARMM General Force Field (CGenFF) version 4.6^19^. Initial parameters for F_430_ with no Ni(I) ion were obtained in a similar manner. Lennard-Jones (LJ) parameters for Ni(I) were fit to water interaction energies from quantum mechanical (QM) calculations. A rigid optimization of Ni(I) with a water molecule (held in TIP3P geometry) was performed to identify the distance of optimal interaction between the ion and water. This optimization was performed with a B971/cc-pVQZ model chemistry, as suggested for transition elements^20^. The interaction energy between these species was then calculated using the same model chemistry, with counterpoise correction for basis set superposition error^21,22^. All QM calculations were performed in Gaussian09^23^. The final LJ parameters for Ni(I) were R_min_/2 = 0.7811 Å and ε = -0.02 kcal/mol.

To enforce the proper coordination geometry of Ni(I) in the macrocyclic ring, harmonic bond terms were added between Ni(I) and the pyrrole nitrogen atoms with a force constant of 500 kcal/mol•Å^2^ and b_0_ of 2.06 Å. Force constants were set to zero for angles and proper dihedral terms involving Ni(I) and the macrocyclic ring, following the CHARMM force field convention to omit explicit contribution to the forces arising from these interactions when involving metals. Improper dihedrals based on the heme residue were added to enforce planarity of the Ni(I) ion within the macrocyclic ring. Afterwards, we refined macrocyclic internal bonds and angles to reproduce the ring puckering and overall macrocyclic conformation from crystallographic data. The final parameters for F_430_ served as the basis to derive mt-F_430_. Additional parameters for methylthio and mercaptopropanamide group substitutions were obtained through CGenFF in a similar manner and later merged with the F_430_ parameters to create the mt-F_430_ and mpa-F_430_ residues. Topology parameters for all cofactors described in this work are available in the Supporting Information.

### Molecular dynamics system setup

To understand the structural relationship between ANME-1 and *M. acetivorans* MCR complexes when bound to different F_430_ cofactors, we built four simulation systems (**Table 1**). To build system *Ma*-F_430_, the atomic coordinates of *M. acetivorans* MCR bound to F_430_ reported by Nayak et al.^24^ were used. For system ANME-mtF_430_, the coordinates for ANME-1 MCR complex bound to mt-F_430_ were obtained from PDB 3SQG^15^. To build system ANME-F_430_, we simply aligned systems *Ma*-F_430_ and ANME-mtF_430_ and transposed the coordinates of each F_430_ molecule to their corresponding active sites. The preparation of the *Ma*-mtF_430_ and *Ma*-mpaF_430_ systems are described below. Protein topologies were generated using CHARMM36m force field^25^ and specific parameters for non-standard amino acids in the structure, namely N^1^-methylated histidine (MHS), 5-methyl-arginine (AGM), thioglycinethioglycin (GL3), didehydroaspartate (DYA), S-methylcysteine (SMC), and 7-hydroxytryptophan (0AF) were obtained from the work of Croitoru et al.^26^. The N-and C-termini of each MCR protein chain were capped with a protonated amine and a deprotonated carboxylate group, respectively. Protonation states of histidine, aspartate, and glutamate residues were assigned by using the PlayMolecule web server based on pK_a_ calculations^27^ and visually checked to ensure an optimized hydrogen-bonding network (Table S1 and S2). Crystallographic waters were retained for all systems except for two water molecules modeled between CoM and CoB in both active sites in the crystal structure reported by Nayak et al., for reasons described below. Methyl groups of CH_3_-S-CoM in both active sites were rebuilt using CHARMM internal coordinate builder^28^. Each enzyme:cofactor complex was solvated in a cubic box of CHARMM-modified TIP3P water^29–31^ and ∼150 mM KCl, including necessary counter ions.

**Table 1.** Composition of each system in this work.

| System ID | Organism | F<sub>430</sub> cofactor | Ions (K<sup>+</sup>/Cl<sup>-</sup>) | # of Waters |
| --- | --- | --- | --- | --- |
| *Ma*-F<sub>430</sub> | *M. acetivorans* | F<sub>430</sub> | 304/242 | 258030 |
| *Ma*-mtF<sub>430</sub> | *M. acetivorans* | mt-F<sub>430</sub> | 304/242 | 258054 |
| *Ma*-mpaF<sub>430</sub> | *M. acetivorans* | mpa-F<sub>430</sub> | 304/242 | 258099 |
| ANME-F<sub>430</sub> | ANME-1 | F<sub>430</sub> | 291/235 | 249030 |
| ANME-mtF<sub>430</sub> | ANME-1 | mt-F<sub>430</sub> | 291/235 | 249081 |

The CHARMM package^28^ was used to perform energy minimizations using the steepest descent algorithm for 100 steps, followed by 500 steps of adopted-basis Newton-Raphson minimization to relax steric clashes. Equilibration of solvent and bulk ions was performed via three independent replicates for 1 ns using OpenMM^32^ version 7.7 with a position restraint force constant of 800 kJ/(mol•nm^2^) on all heavy atoms, including those of the cofactors. A 2-fs integration step was used with a Langevin integrator and a temperature of 310 K, while a Monte Carlo barostat algorithm was applied to maintain the pressure at 1 bar. Short-range van der Waals forces were switched smoothly to zero from 10-12 Å and electrostatic interactions were calculated via the Particle Mesh Ewald method^33,34^ with a real-space cutoff of 12 Å. The equilibrated system coordinates of all 3 replicates were submitted to unrestrained production runs of 100 ns each, while saving coordinates every 10 ps.

As mentioned before, the presence of Val419 in the α subunits of ANME-1 MCR accommodates the methylthio group of mt-F ^15^, whereas methanogen MCRs possess a glutamine in the same position that could result in steric clashes with the modifications at the 17^2^ position of F_430_ – i.e. the methylthio group in mt-F_430_ or the mercaptopropanamide group in mpa-F_430_. To study possible active-site rearrangements necessary to properly fit 17^2^-modified F_430_s into the *M. acetivorans* MCR binding site, we built system *Ma-*mtF_430_ and *Ma-*mpaF_430_ by applying an alchemical transformation protocol: electrostatic and Lennard-Jones potentials of the modified F_430_ cofactors were switched from 0 (no interaction) to 1 (full interaction) in a stepwise manner through a λ scaling factor applied to the Hamiltonian^35,36^. Lennard-Jones potentials were switched on first (λ=0.00, 0.05, 0.10,0.15, 0.20, 0.30, …, 0.70, 0.75, 0.80, 0.85, 0.90, 0.95, 1.00) and electrostatic interactions were switched on afterwards (λ=0.00, 0.10, 0.20, …, 0.80, 0.90, 1.00). Such an approach allows for a gentle adjustment of the protein binding site residues. We built the system using CHARMM as described above. Before energy minimization, the system’s coordinates were used to rebuild their topologies in GROMACS using the *pdb2gmx* utility. The cofactor force field parameters were converted to GROMACS format using the *cgenff_charmm2gmx.py* script available at https://github.com/Lemkul-Lab/cgenff_charmm2gmx. The alchemical transformation protocol in each λ window consisted of a) energy minimization using steepest descent algorithm for 1000 steps, followed by another energy minimization using the conjugate gradient algorithm for 1000 steps; b) an equilibration under an NPT ensemble using position restraints on all heavy atoms [force constant of 800 kJ/(mol•nm^2^)] for 100 ps using the nonbonded interaction parameters described above and c) an unrestrained production run for 1 ns. Soft-core α and σ parameters were set to 0.5 and 0.3, respectively. ∂H/∂λ and ΔH values were written every 0.02 ps. Throughout the protocol, we applied flat-bottom restraints as shown in **Figure S1** to preserve the overall orientation of the F_430_ cofactors relative to their respective binding sites. The initial restraining force constant of 200 kJ/(mol•nm^2^) was decreased linearly as the λ value increased such that the force constant was zero when λ=1. The last frame of the production run in each λ window was used as input for the minimization step of the next λ window. At the completion of the alchemical transformation, the last frame of the λ=1 production run was used to initiate three independent replicas following the 1-ns equilibration and 100-ns production protocols used for the other systems.

### Electric field calculations

To understand the role of electrostatic forces on methane formation by MCR, we used TUPÃ^37^ to calculate the electric fields organized by the different MCR complexes acting on the methyl-sulfur bond of CH_3_-S-CoM to facilitate its cleavage. We used the BOND mode and defined the bond vector as S → CH_3_ such that any field vector aligned with the bond vector results in positive values. All protein, F_430_, and HS-CoB atoms were included in the environment set (*i.e.*, the atoms exerting the electric field) while self-contribution of atoms in CH_3_-S-CoM were ignored. The complete configuration parameters used in the calculation can be found in Supporting Information.

## Results and Discussion

### Initial simulations to establish a starting structure

The majority of MCR structures solved by X-ray crystallography and available in the PDB, such as entries 5N1Q, 1E6Y, 3POT, 3M2R and 5A0Y^38–42^, show an electron density between CoM and CoB. In many of these structures, this electron density is modeled as a water molecule trapped inside a hydrophobic cage composed by Phe and Tyr residues surrounding CoM. Considering that the reconstruction of the methyl group of CH_3_-S-CoM caused some steric clash with that water molecule, we decided to check if the presence of water in this location had any structural impact in the active site. To do so, we built a system with *M. acetivorans* MCR^24^ that contains the canonical F_430_ (*Ma-*F_430_) including these two water molecules and tracked their dynamics during equilibration and production runs. As shown in Movie S1, the water molecule remained trapped inside the hydrophobic cage throughout equilibration due to the restrained motions of protein atoms, but it quickly left the active site once the restraints were released.

As shown in Figure 2, the trapped water molecules left both active sites in all replicates shortly after the unrestrained production run begins. Throughout the 100 ns of simulation in each replicate, no water molecule was able to diffuse inside the hydrophobic cage or return to its original position as modeled in the crystal structure. These results suggest that a water molecule cannot be located between CH_3_-S-CoM and HS-CoB, though the structure obtained via X-ray crystallography contains HS-CoM, not its methylated form. It is also possible that, given the lack of electronic polarization of our model (and thus the inability of water molecules to depolarize in response to their surrounding microenvironments), it is expected that these water molecules in the active site would be repelled in our simulations. However, the strongly hydrophobic nature of the microenvironment suggests that such repulsion would occur even when accounting for more electronically robust models. Further, all structures were experimentally obtained with F_430_ cofactors with Ni(II), which are chemically distinct from the Ni(I) state used in our simulations. Nevertheless, we could not provide rationale for possible structural roles a water molecule could have in MCR pre-catalytic active site. Further studies should be carried to more confidently determine if the observed electron densities do represent water molecules and, if so, what kind of role they have in the active site, or if such electron densities could represent other apolar species with a similar number of electrons (e.g. CH_4_).

**Figure 2.**
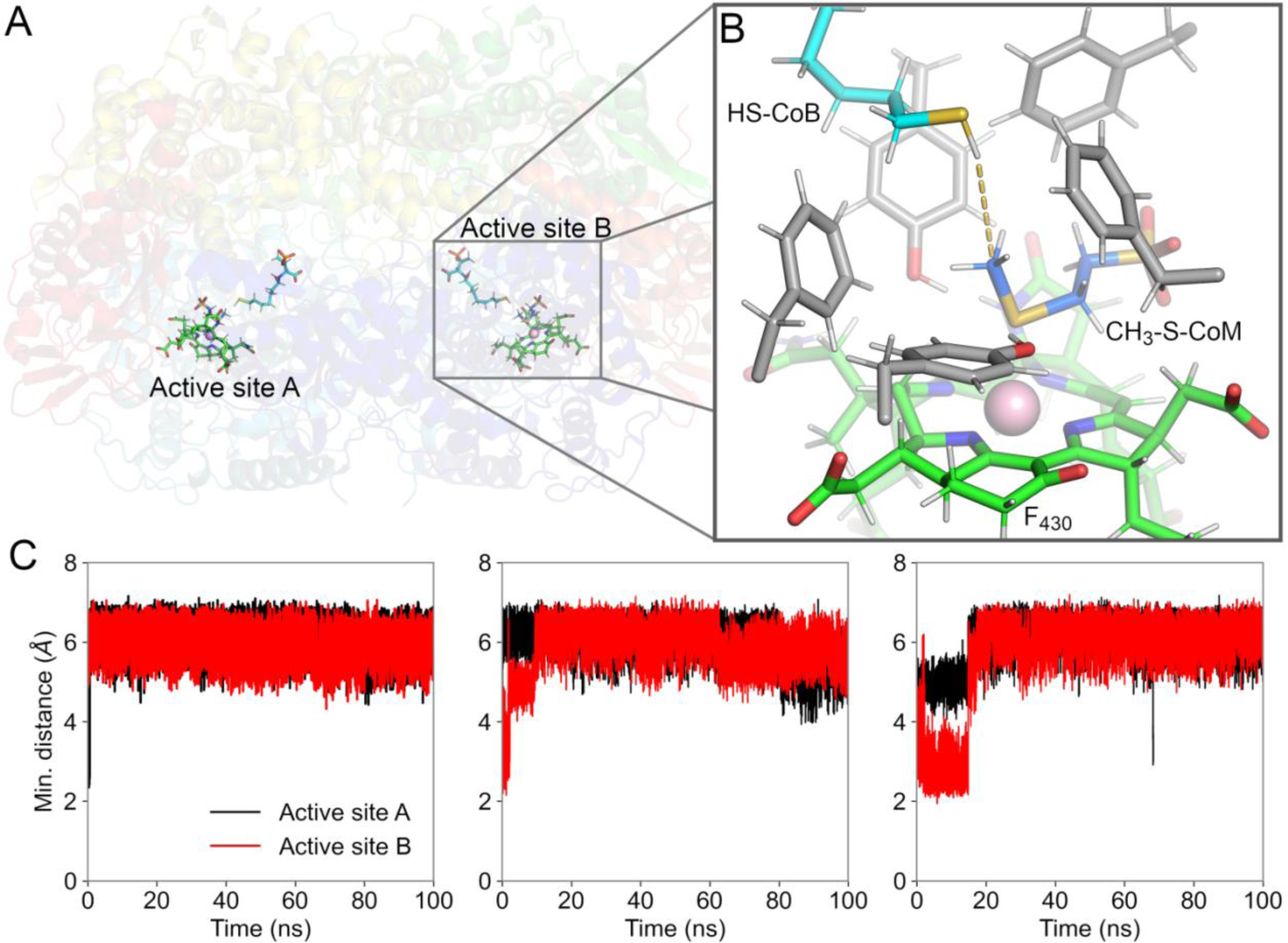
Hydrophobic cage architecture in the MCR active site. A) Overall structure of *M. acetivorans* MCR bound to F_430._ Active sites A and B are shown. B) Aromatic and hydrophobic residues composing the hydrophobic cage are shown in light gray, shielding the reaction microenvironment from water molecules. HS-CoB are shown in cyan; CH_3_-S-CoM are shown in blue and F_430_ cofactors are shown in green. C) Minimum distances between the methylthio group of CH_3_-S-CoM molecules in active sites A and B and any water molecule in the system.

### Active site structure and dynamics

To explore the structural features adopted by MCRs from *M. acetivorans* compared to ANME-1, we evaluated the active site dynamics of the two enzymes in the presence of the canonical F_430_ compared to mt-F_430_ (Figure 1A). Recently, we have identified a modified F_430_ in *M. acetivorans* with a structure that we preliminarily assign as 17^2^-mercaptopropanamide-F_430_ (mpa-F_430_) (see Supplemental Information). Thus, simulations were additionally performed with this extended mercaptopropanamide modification. To evaluate the dynamics in all systems and their impact on the coordination among F_430_, CH_3_-S-CoM and HS-CoB, we calculated key distances that characterize the coordination between these molecules in the active site (Figure 3). First, we defined D1 as the distance between the methyl carbon of CH_3_-S-CoM and the sulfur atom of HS-CoB. We also defined D2 as the distance between the Ni(I) of F_430_ cofactor and the thioether sulfur of CH_3_-S-CoM. Given the symmetry between active sites A and B, we pooled the data of both sites for each system and analyzed them together.

**Figure 3.**
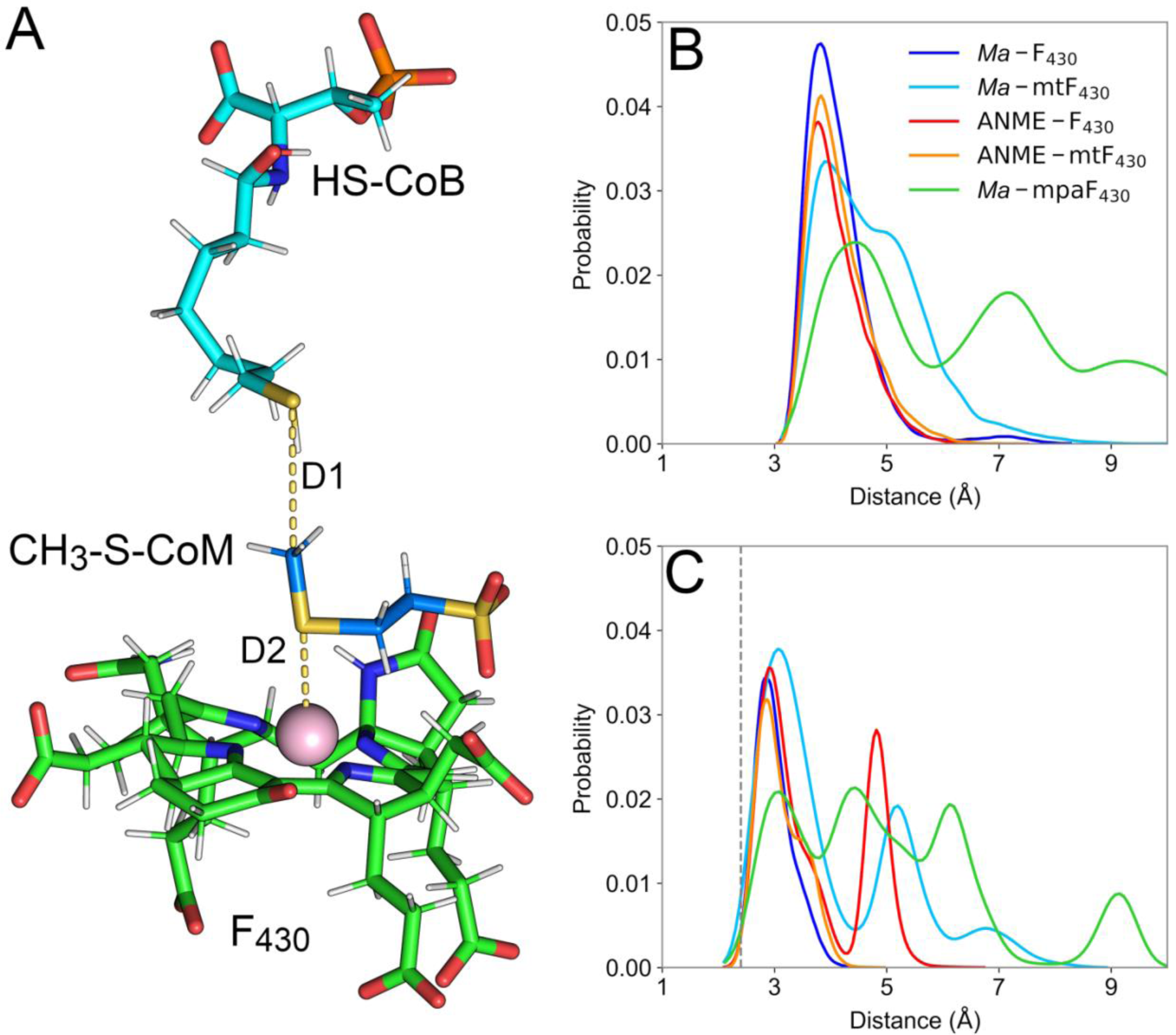
Overall cofactor and substrate coordination. A) Pre-reactant scheme of MCR bound to F_430_, CH_3_-S-CoM, and HS-CoB. Key distances were designated as D1 and D2, representing the distance between the methyl group of CH_3_-S-CoM and the thiol sulfur atom of HS-CoB and the distance between Ni(I) of F_430_ cofactors and methylthio sulfur atom of CH_3_-S-CoM, respectively. B) Distribution probability of distance D1 for each of the enzyme:cofactor complexes studied here. C) Distribution probability of distance D2 for the same systems. The gray dashed line represents the crystallographic distance, for reference.

For all replicates of the ANME*-*mtF_430_ and *Ma-*F_430_ systems, D1 and D2 were consistent throughout the trajectories and coordination among cofactors were well maintained. For ANME*-*mtF_430_, D1 and D2 averaged at 4.2 ± 0.5 Å and 3.1 ± 0.4 Å, respectively. For *Ma-*F_430_, D1 and D2 averaged at 4.1 ± 0.5 Å and 3.7 ± 0.9 Å, respectively. It is important to note that the reference crystal structures for both systems were obtained with HS-CoM present in the active site, which yields a smaller distance for D2 of 2.4 Å, likely due to the different chemical nature of the interaction between Ni(II) and a thiol compared to Ni(I) and a thioether.

The presence of Val419 in the α subunit of ANME-1 MCR creates a pocket that accommodates the F_430_ modification in the 17^2^ position^15^. To test the impact of the methylthio group of mt-F_430_ in maintaining ANME-1 active site structure, we simulated ANME-1 MCR bound to unmodified F_430_. In our ANME*-*F_430_ simulations, some replicates maintained the canonical coordination between F_430_ and CH_3_-S-CoM. D1 had an average of 4.1 ± 0.5 Å across all replicates, but D2 varied among replicate simulations and active sites. For the first replicate, D2 averaged 3.1 ± 0.4 Å in active site A and 3.2 ± 0.4 Å in active site B. For the second and third replicates, D2 increased in one of the active sites (average of 4.8 ± 0.5 Å) while maintaining an average of 3.0 ± 0.4 Å (Figure 4A) at the other active site. Upon further investigation, we were able to correlate the increase in D2 with events in which HS-CoB diffused further into the active site, forcing a dihedral rotation in CH_3_-S-CoM that disrupted the surrounding hydrophobic cage structure, consequently increasing the distance between CH_3_-S-CoM and F_430_ core (Figure 4B).

**Figure 4.**
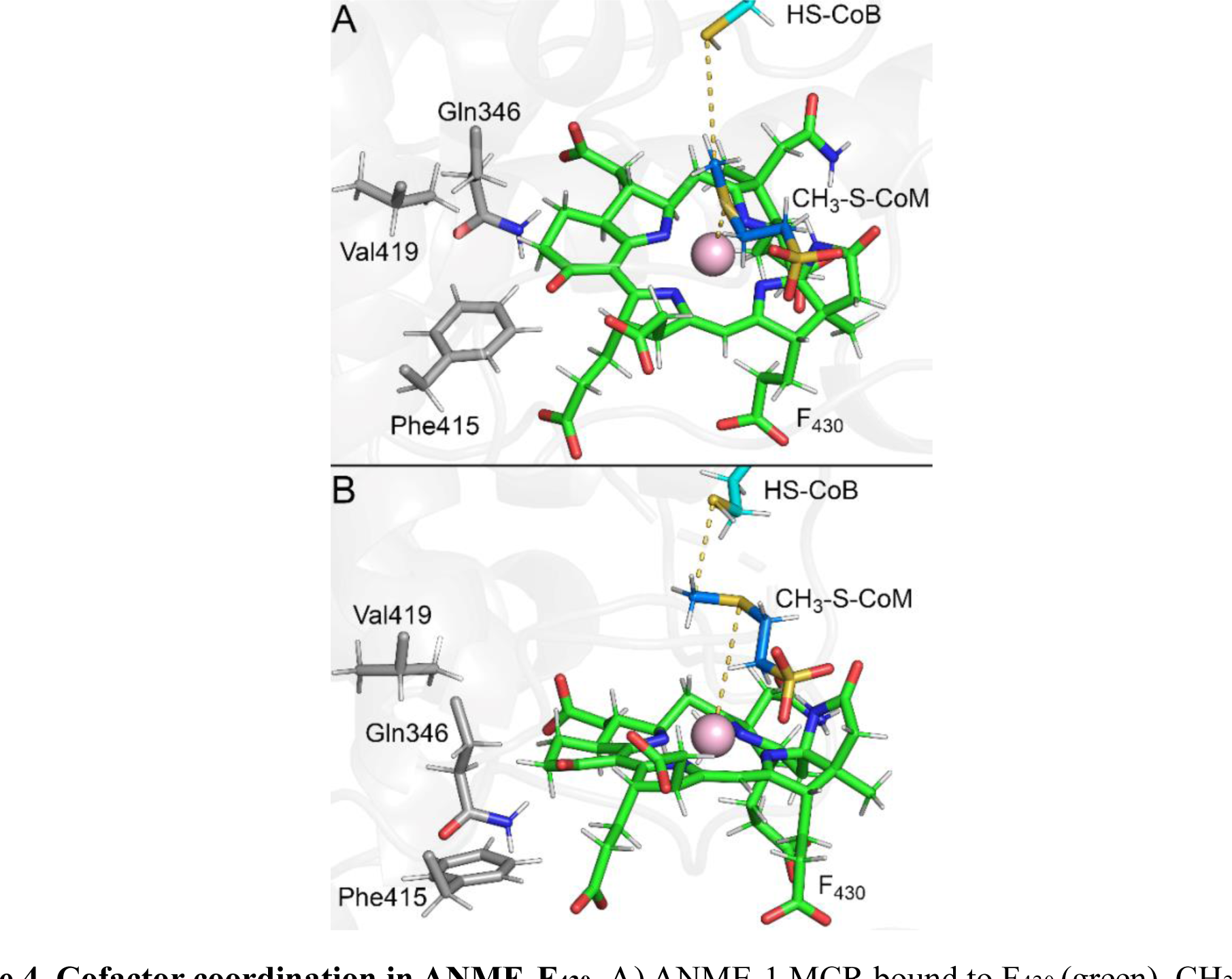
Cofactor coordination in ANME*-*F_430_. A) ANME-1 MCR bound to F_430_ (green), CH_3_-S-CoM (blue) and HS-CoB (cyan). Key MCR residues are shown in gray sticks. Representative structure of canonical coordination among cofactors observed in our simulations. B) Representative structure of the increased D2 between CH_3_-S-CoM and F_430_.

In *M. acetivorans* MCR, Gln420 occupies the same site as Val419 in ANME-1 MCR, which is expected to create a steric clash with F_430_ cofactors containing a modification at the 17^2^ position. In our simulations of *Ma-*mpaF_430_ and *Ma-*mtF_430_ systems, the F_430_ modifications in the 17^2^ position impacted the coordination between cofactors. In most replicates of *Ma-*mpaF_430_, the large mercaptopropanamide group clashed with Gln420 and interacted with Phe416, which destabilized the coordination between CH_3_-S-CoM and mpa-F_430_ (Figure 5A). In one replicate, however, the mercaptopropanamide group was able to dislocate Gln420 and accommodate itself between Phe416 and Gln345 (Figure 5B), thus maintaining the canonical coordination among the cofactors throughout the simulation. In another replicate in which the mercaptopropanamide group adopted an extended conformation, the canonical coordination between CH_3_-S-CoM and mpa*-*F_430_ through the thioether sulfur -Ni(I) interaction was broken, and through a rearrangement, CH_3_-S-CoM interacted with mpa-F_430_ via its sulfonate group for over 80 ns, somewhat similar to the pose required in the reaction mechanism proposed by Patwardhan *et. al.*^11^. Such an alternative pose was observed in only one active site and yielded D1 and D2 distances of 4.4 ± 0.7 Å and 5.8 ± 0.7 Å, respectively, while the average minimum distance between the sulfonate oxygen atoms and the Ni(I) of mpa-F_430_ was 1.76 ± 0.08 Å (Figure 5C).

**Figure 5.**
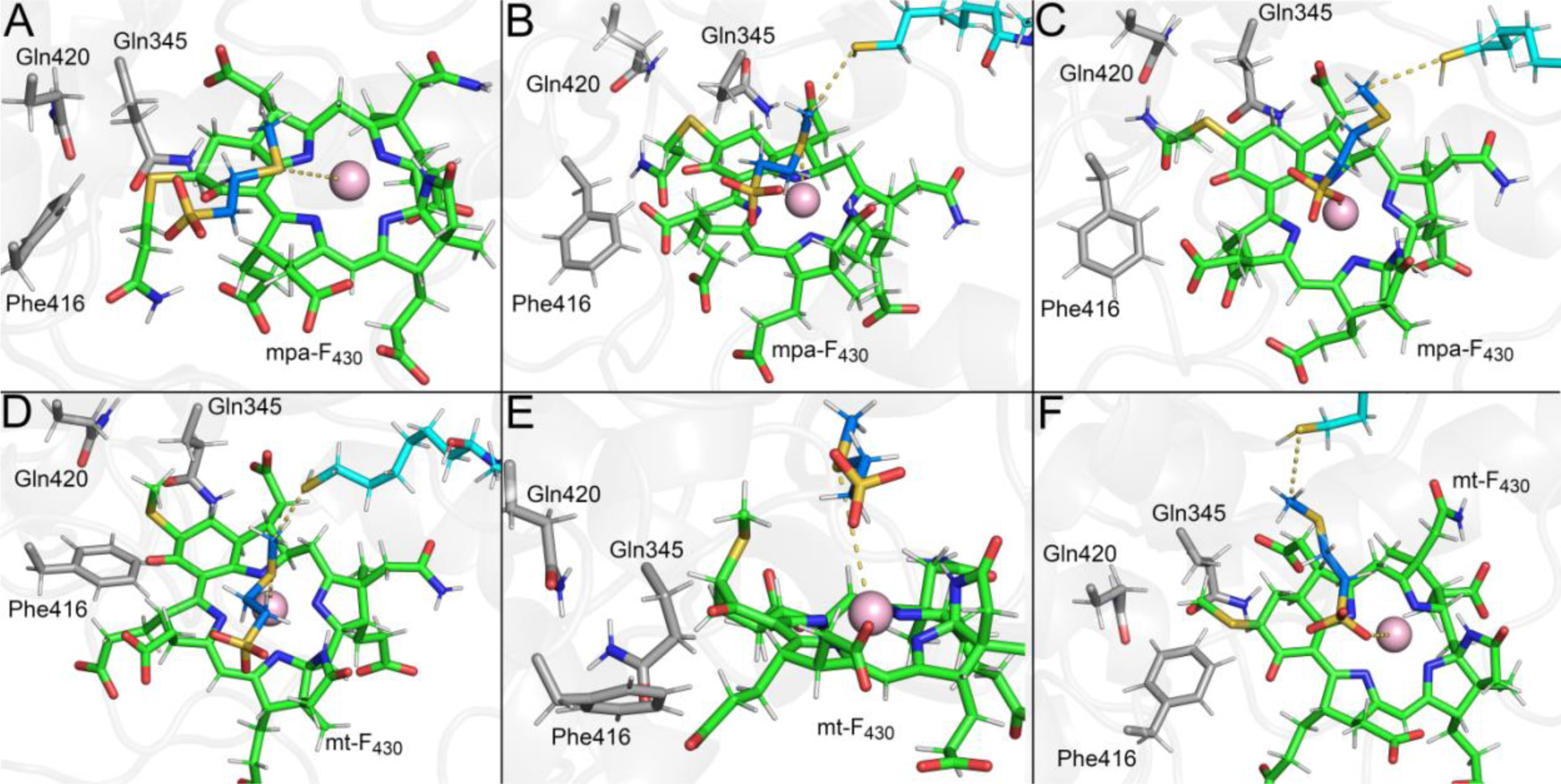
Cofactor coordination in *Ma-*mpaF_430_ and *Ma-*mtF_430_. A-C) Representative structures of *M. acetivorans* MCR bound to mpa-F_430_ (green), CH_3_-S-CoM (blue) and HS-CoB (cyan) sampled in our simulations. Key MCR residues are shown in gray sticks. D-F) Representative structures of *M. acetivorans* MCR bound to mt-F_430_ (green).

For *Ma-*mtF_430_, the methylthio group of mt-F_430_ also suffered from steric clashes from Gln420, but to a lesser extent, which was accommodated by reorganization within the active site in some cases. In the replicates in which the mt-F_430_ methylthio group was able to be better accommodated in the active site, D1 and D2 averaged 3.4 ± 0.5 Å and 4.6 ± 0.8 Å, respectively, maintaining the canonical coordination among the cofactors. In such cases, Phe416 and Gln345 maintained their original interactions with mtF_430_, while Gln420 repositioned to better accommodate the methylthio group (Figure 5D). In the replicates in which the coordination was lost, Gln345 repositioned within the active site, leading to a change in the mtF_430_ 6-member ring puckering, leading the methylthio group to interact with the methylated end of CH_3_-S-CoM, destabilizing their coordination with the mt-F_430_ core (Figure 5E). Analogous to the alternative pose observed in *Ma-*mpaF_430_, one replicate of our *Ma-*mtF_430_ simulations sampled a configuration in which the sulfonate group of CH_3_-S-CoM was not interacting with Ni(I) of mt-F_430_ directly (average minimum distance between Ni(I) and the sulfonate oxygens was 3.9 ± 0.2 Å), but the overall configuration resembled the alternative pose observed in *Ma-*mpaF_430_ (Figure 5C). In that pose, D1 and D2 average distances were 4.5 ± 0.6 Å and 6.8 ± 0.6 Å, respectively.

The observed accommodation (or lack thereof) of different F_430_ cofactors suggests a delicate balance among the amino acids of the MCR enzyme and the cofactor and substrates. Important among these residues are those in the hydrophobic cages. The presence of conserved Phe and Tyr residues in the hydrophobic cages surrounding the MCR active sites suggests the importance of aromatic character within the active site microenvironment for the reaction being catalyzed. To better understand the organization of such structures throughout our simulations of each enzyme:cofactor system, we calculated the average of the distances between the aromatic residues composing the cage and the methyl group of CH_3_-S-CoM, from now on referred as D3_avg_ (Figure 6A**,B**). For *M. acetivorans*, these residues were defined as Phe343, Tyr346, and Phe463 of subunit α, and Phe359 and Tyr365 of subunit β. In ANME-1, residues Phe344, Tyr347, and Phe462 of subunit α, and Phe357 and Tyr363 of subunit β were used.

**Figure 6.**
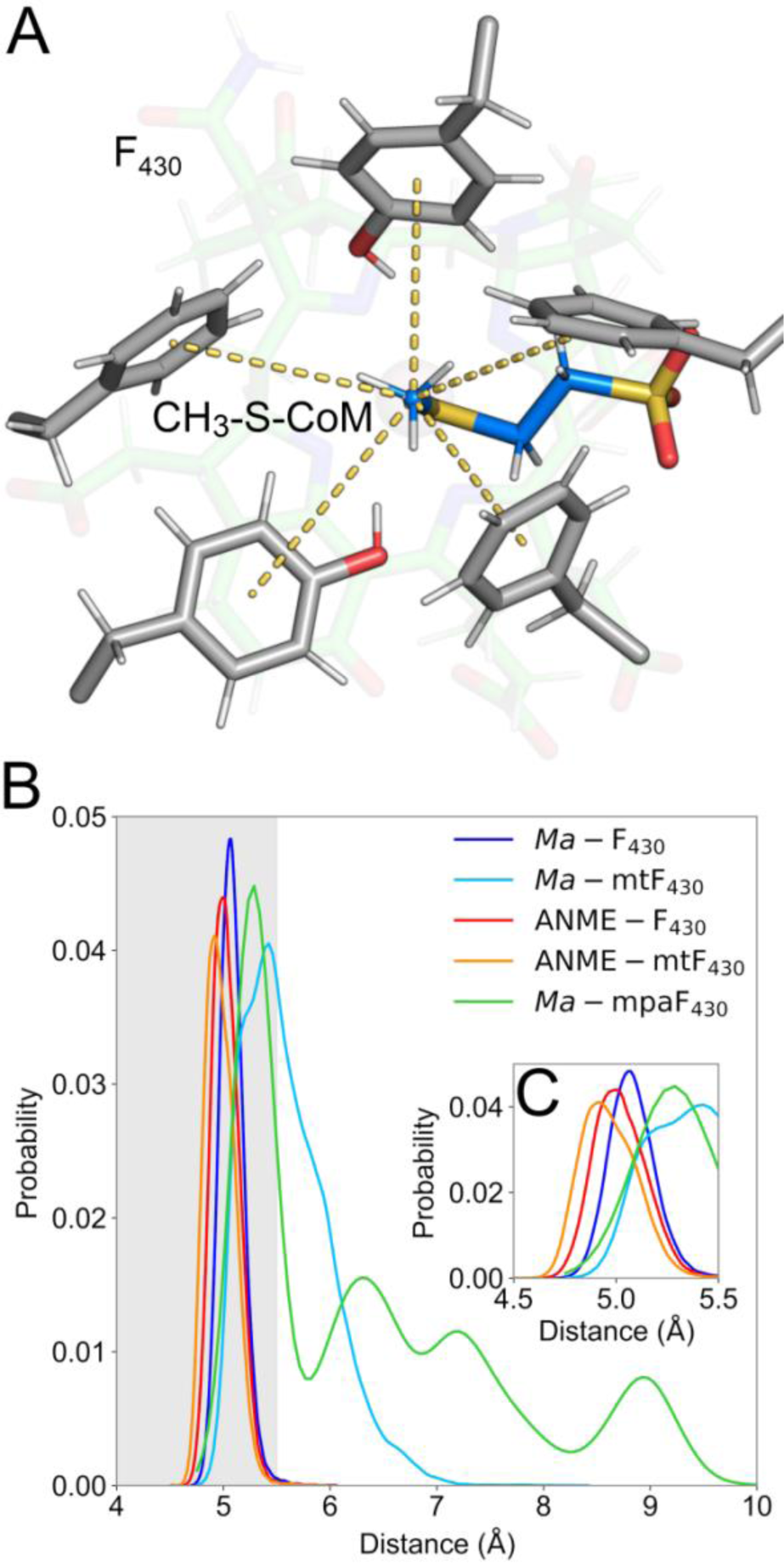
Overall organization of hydrophobic cage. A) Pre-reactant scheme of MCR bound to F_430_, CH_3_-S-CoM, and HS-CoB. Conserved residues comprising the hydrophobic cage are shown in gray and their distances to the thioether methyl group of CH_3_-S-CoM are highlighted in yellow. B) Probability distribution of D3_avg_ composed by distances highlighted in panel A for each of the enzyme:cofactor complexes studied here. Gray areas represent commonly observed distances for hydrophobic interactions. C) Highlight of the distributions between 4.5 Å and 5.5 Å.

For systems *Ma-*F_430_, ANME*-*mtF_430_ and ANME-F_430_, D3_avg_ were 5.1 ± 0.1 Å, 5.0 ± 0.1 Å and 5.0 ± 0.1 Å, respectively. As shown in Figure 6C, ANME*-*mtF_430_ and ANME-F_430_ sampled lower D3_avg_ values slightly more often in comparison to *Ma-*F_430_. Not surprisingly, these are the systems in which the canonical coordination among cofactors is better maintained throughout the simulations, suggesting that such hydrophobic cage also plays a structural role in maintaining the overall active site configuration. For systems *Ma-*mtF_430_ and *Ma-*mpaF_430_, in which the canonical coordination among cofactors was less frequently observed, D3_avg_ were 5.6 ± 0.4 Å and 6 ± 1 Å, respectively, with a broad distribution in longer distance values. These results emphasize the correlation between the loss of canonical coordination between cofactors and the cage structural organization, and stress the structural impact of the steric clash between F_430_ modifications and active site residues, specially Gln420 of subunit α.

### MCR electric fields driving catalysis

The superior affinity of an enzyme to its substrate’s transition state is derived from both conformational and electronic properties^43,44^. The role of intrinsic electric fields driving enzyme-mediated catalysis was proposed by Warshel in the 1970s^43,45,46^ and gained traction more recently with the development of more robust computational models and experimental techniques to measure such effects^44,47–49^. Electric fields exerted by enzymes act at specific chemical bonds of the substrate, lowering the reaction activation barrier and stabilizing the transition state.

For the methane-formation reaction catalyzed by MCR, the currently accepted mechanism involves one-electron chemistry such that the thioether bond of CH_3_-S-CoM is homolytically cleaved, forming a methyl radical and a covalent bond between Ni(II) and S-CoM. To understand how MCR organizes its active site electronic environment to drive the methane-forming reaction, we calculated the total electric field (^𝐸⃗^ _𝑡𝑜𝑡𝑎𝑙_) acting at the center of thioether S-CH_3_ bond of CH_3_-S-CoM (Figure 7A) for each enzyme:cofactor system. From ^𝐸⃗^ _𝑡𝑜𝑡𝑎𝑙_, we calculated the effective electric field (^𝐸⃗^ _𝑒𝑓𝑓_), which is the vectorial projection of ^𝐸⃗^ _𝑡𝑜𝑡𝑎𝑙_ onto the bond and represents the field that is effectively modulating the bond dipole. Finally, we also calculated an angle, χ, between the ^𝐸⃗^ _𝑡𝑜𝑡𝑎𝑙_ and ^𝐸⃗^ _𝑒𝑓𝑓_ field vectors as a measure of their alignment (Figure 7A, inset). That is, the value of χ allows us to determine how efficient the total electric field is in promoting the desired chemical reaction. To allow cross comparison between different systems, we only considered the active sites and replicates in which the canonical coordination among the cofactors were reasonably maintained.

**Figure 7.**
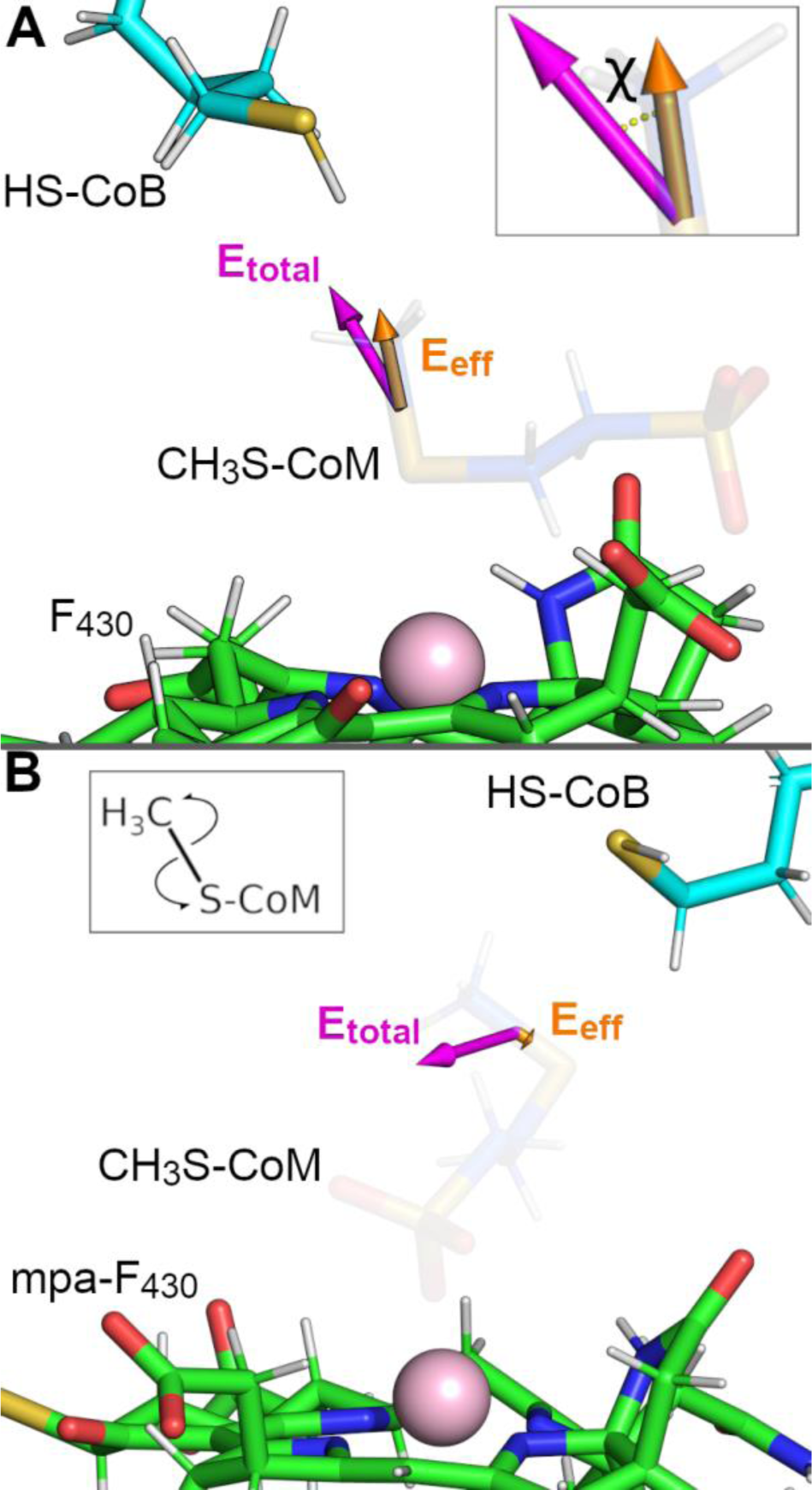
Electric field organization driving MCR catalysis. A) Total and effective electric fields (magenta and orange, respectively) exerted by MCR, HS-CoB, and F_430_ acting at the thioether S-CH_3_ bond of CH_3_-S-CoM. Inset depicting the angle χ measuring the alignment between ^𝐸⃗^ _𝑡𝑜𝑡𝑎𝑙_ and E_eff_ field vectors. B) Total and effective electric fields (magenta and orange, respectively) exerted by MCR, HS-CoB and mpa-F_430_ acting at the thioether S-CH_3_ bond of CH_3_-S-CoM considering its alternative pose. Inset depicting a scheme of the homolytic thioether bond cleavage.

As represented in Figure 7A, ^𝐸⃗^ _𝑡𝑜𝑡𝑎𝑙_ of all systems were mainly positive, meaning they were oriented in the same direction as the bond dipole vector (S → CH_3_). The good alignment between ^𝐸⃗^ _𝑡𝑜𝑡𝑎𝑙_ and the bond implies that such a field is able to polarize the bond dipole even further and facilitate bond cleavage. As shown in **Table 2**, the average ^𝐸⃗^ _𝑡𝑜𝑡𝑎𝑙_ for *Ma-*F_430_, *Ma-* mtF_430_ and *Ma-*mpaF_430_ were 39 ± 13, 34 ± 15, and 19 ± 12 MV/cm, respectively. For ANME*-* mtF_430_ and ANME-F_430_, the average ^𝐸⃗^ _𝑡𝑜𝑡𝑎𝑙_ were 67 ± 13 and 58 ± 12 MV/cm, respectively.

**Table 2.** MCR intrinsic electric field values acting at thioether bond of CH_3_-S-CoM.

| System | Average $\vec{E}_{total}$ (MV/cm) | Average $\vec{E}_{eff}$ (MV/cm) | Alignment Angle $\chi$ (°) |
| --- | --- | --- | --- |
| *ma*-F<sub>430</sub> | $52 \pm 13$ | $39 \pm 13$ | $39 \pm 16$ |
| *ma*-mtF<sub>430</sub> | $55 \pm 13$ | $34 \pm 15$ | $42 \pm 21$ |
| *ma*-mpaF<sub>430</sub> | $54 \pm 14$ | $19 \pm 12$ | $69 \pm 13$ |
| *anme*-mtF<sub>430</sub> | $67 \pm 13$ | $56 \pm 14$ | $30 \pm 16$ |
| *anme*-F<sub>430</sub> | $58 \pm 12$ | $50 \pm 13$ | $28 \pm 14$ |

Except for ANME-mtF_430_, the average ^𝐸⃗^ _𝑡𝑜𝑡𝑎𝑙_ for all systems was close to ∼55 MV/cm, which can be considered weak in comparison to previous reports for other enzymes^47,50^. This property may be related to the proposed homolytic bond cleavage as opposed to heterolytic bond cleavage.^6^ For ^𝐸⃗^ _𝑒𝑓𝑓_, ANME-mtF_430_ and ANME-F_430_ presented higher averages, specifically measuring 56 ± 14 and 50 ± 13 MV/cm, respectively, whereas *Ma-*F_430_, *Ma-*mtF_430_, and *Ma-* mpaF_430_ produced ^𝐸⃗^ _𝑒𝑓𝑓_ averages of 39 ± 13, 34 ± 15 and 19 ± 12 MV/cm, respectively.

The pronounced differences in ^𝐸⃗^ _𝑒𝑓𝑓_ of each system reflect the alignment between the total field and the bond. As shown in **Table 2**, the χ angle differed among systems, suggesting the active site conformation does impact ^𝐸⃗^ _𝑒𝑓𝑓_ . The *Ma-*mpaF_430_ and *Ma-*mtF_430_ systems had the largest χ average angle across the systems, 69 ± 13**°** and 42 ± 21**°**, respectively, implying that these active site conformations are the most perturbed. In fact, even in maintaining the canonical cofactor coordination, mpa-F_430_ had a D3_avg_ of 5.9 ± 0.3 Å, highlighting the extent to which the active site structural organization impacts intrinsic electric fields in MCR. Interestingly, the lowest average χ angle values across the systems were observed for ANME-mtF_430_ and ANME-F_430_, suggesting ANME-1 MCR structured its active site to optimize the electric field acting on the thioether S-CH_3_ bond of CH_3_-S-CoM.

These results are somewhat surprising since ANME-1 catalyzes the reverse reaction (methane oxidation) but both of our systems using ANME-1 yielded stronger ^𝐸⃗^ _𝑒𝑓𝑓_ values in comparison to *M. acetivorans*, suggesting a better catalytic efficiency of the former in comparison to the later. However, given the complex electronic nature of the active site residues and cofactors involved in the reaction, a proper quantitative confirmation of the magnitudes observed in this work should be reassessed by models capturing electronic polarization effects.

Moreover, we evaluated the same electric field organized by MCR but now considering the alternative pose observed for *Ma*-mpaF_430_, and similar to the alternative binding scheme leading to long-range electron transfer steps proposed by Patwardhan *et. al.*^11^ (Figure 7B). Throughout the trajectory, the methyl group of CH_3_-S-CoM presented an increased degree of flexibility, often rotating about the thioether bonds while still maintaining the interaction between CH_3_-S -CoM and mpa-F_430_ through its sulfonate group. In most frames, ^𝐸⃗^ _𝑡𝑜𝑡𝑎𝑙_ was poorly aligned with the thioether S-CH_3_ bond (average χ = 64 ± 20°), often sampling electric fields nearly perpendicular to the bond, resulting in a very weak ^𝐸⃗^ _𝑒𝑓𝑓_ (average of 16 ± 12 MV/cm). At the same time, the structure of the hydrophobic cage residues surrounding CH_3_-S-CoM was very poorly maintained in this replicate, with a D3_avg_ of 6.5 ± 0.6 Å. Given the weak ^𝐸⃗^ _𝑒𝑓𝑓_ acting on the thioether bond in the frames sampling the alternative pose of CH_3_-S-CoM within the active site, this alternative pose is not expected to support catalysis in the context of intrinsic electric fields organized by MCR. However, since our analyses reported here show that the 17^2^ F_430_ modifications result in large perturbations to the *M. acetivorans* MCR active site, other aspects of the reorganization may be responsible for the impact on the measured electric fields. Additionally, we recognize such fields should be recalculated with models with more robust electronic description to account for the intricate electronic properties of the active site microenvironment.

To further decompose the contributions of different active-site residues to ^𝐸⃗^ _𝑒𝑓𝑓_ in each system, we calculated the contribution of only the hydrophobic cage residues, finding that they can be responsible for up to half of the magnitude of ^𝐸⃗^ _𝑒𝑓𝑓_ . As shown in **Table 3**, such residues contributed 12 ± 11, 20 ± 13, and 6 ± 10 MV/cm for systems *Ma-*F_430_, *Ma-*mtF_430_, and *Ma-*mpaF_430_, respectively, while analogous hydrophobic cage residues of ANME-1 contributed 24 ± 13 and 21 ± 11 MV/cm for systems ANME-mtF_430_ and ANME-F_430_, respectively. These results correlate with the D3_avg_ distances of shown in Figure 6C in which ANME-mtF_430_ and ANME-F_430_ sampled shorter distances more frequently than other systems. These results suggest the hydrophobic cage residues not only play a structural role in maintaining the canonical coordination among cofactors but also play an electronic role driving methane formation catalysis.

**Table 3.**
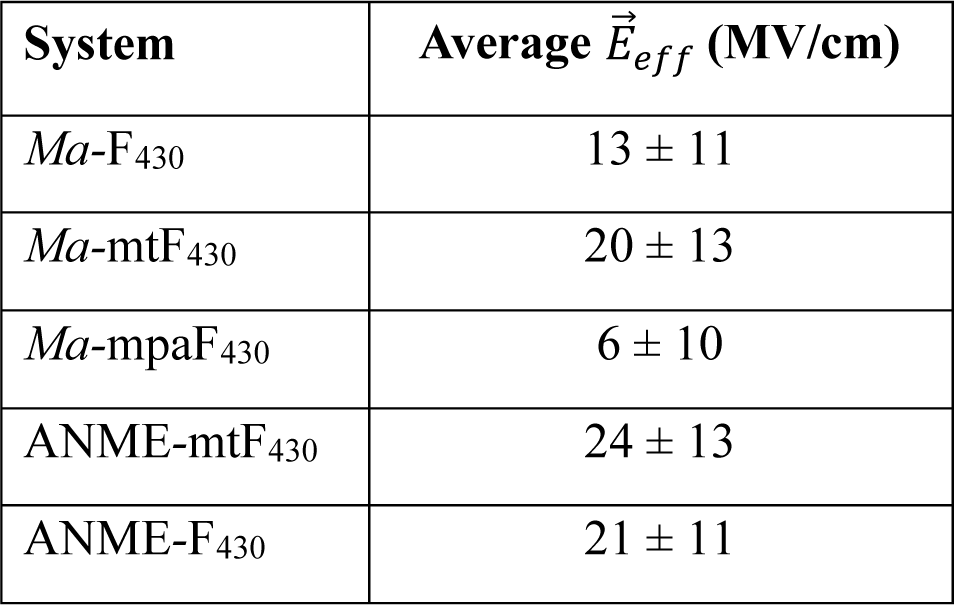
Contribution of hydrophobic cage residues to MCR intrinsic electric field values acting at thioether bond of CH_3_-S-CoM.

## Conclusions

Through our MD simulations, we provided key insights into MCR dynamics from both *M. acetivorans* and ANME-1 when bound to modified F_430_ cofactors and substrates for the methane formation reaction. First, we investigated the presence of electron densities inside the active site commonly modeled as water molecules in X-ray structures. Our simulations suggest that the presence of waters inside a hydrophobic cage composed by aromatic residues surrounding CH_3_-S-CoM is unlikely to be favorable, given the chemical nature of the active site and the rapid expulsion of water molecules in all our simulations. Our observations indicate that future studies are still necessary to determine the chemical nature of these observed electron densities and the potential significance of water molecules in the active site.

Our simulations enabled us to characterize the dynamics of enzyme:cofactor complexes with F_430_ cofactors containing modifications in the 17^2^ position and their relationship with active site dynamics and coordination among cofactors. We observed an absolute prevalence of the active site structure organization for all replicates of *Ma-*F_430_ and ANME-mtF_430_, suggesting our model is able to reasonably capture the active site structure and canonical cofactor coordination. Moreover, we demonstrated the impact of the steric clash between Gln420 of *M. acetivorans* MCR subunit α and the 17^2^ methylthio and 3-mercaptopropanamide F_430_ modifications on the active site structure. The presence of such bulkier groups perturbed the active site interaction network and impacted the structural organization of the hydrophobic cage residues surrounding CH_3_-S-CoM, in turn leading to the loss of the canonical coordination among cofactors representing the pre-catalytic state. However, we did observe some evidence of active site reorganization that better maintained the canonical configuration and suggests that modifications could potentially be accommodated. Interestingly, we also observed an alternate binding pose in some replicates of *M. acetivorans* MCR with mpa-F_430_ and mt-F_430_ in which CoM interacts with Ni(I) through its sulfonate group instead of the thioether sulfur. In ANME-1, Val419 of subunit α occupies the analogous position as Gln420 of *M. acetivorans*, creating a suitable pocket to accommodate the 17^2^ methylthio group of mt-F_430_. In simulations of ANME-F_430_, the absence of the methylthio group also impacted the active site structure and led to some loss of the canonical cofactor coordination, although to a lesser extent when compared to systems *Ma-*mtF_430_ and *Ma-*mpaF_430_.

Finally, we characterized the intrinsic electric fields organized by MCR in its active sites to drive the methane formation reaction. We showed that MCR exerts an electric field aligned with the thioether bond of CH_3_-S-CoM, which polarizes the bond dipole and facilitates the bond homolytic cleavage and the formation of the methyl radical. Such an electric field is in line with the prevailing hypothesis for the mechanism through this reaction proceeds. Moreover, we also demonstrated how these fields are sensitive to the hydrophobic cage structural organization. In the replicates in which the structure was perturbed, the total electric field was weaker and misaligned with the bond. Yet, due to the complex electronic nature of the active site molecular components, the electric fields studied here should be revisited by future efforts employing electronically polarizable models to quantitatively capture intricate electronic properties of MCR active sites.

Overall, our results contribute to a better understanding of MCR active site dynamics and the impact of F_430_ modifications, as well as how the enzyme organizes intrinsic electric fields to drive catalysis and the role of the active site structure in the maintenance of such fields.

## Funding

This work was supported by the US Department of Energy (DOE), Office of Science, Basic Energy Sciences (BES) (Grant DE-SC0022338).

## Acknowledgments

We thank Kaleb Boswinkle and Aleksei Gendron for their work on identifying the proposed mpa-F_430_ in *M. acetivorans*. We also thank Satish Nair and William W. Metcalf for providing the coordinates of *M. acetivorans* MCR, and Virginia Tech Advanced Research Computing for computing time and resources.

## Supporting Information Available

Additional methodological details for the characterization of mpa-F_430_ and the flat-bottom restraint used during alchemical transformations, two additional figures illustrating the flat-bottom restraint scheme and the mass spectra and UV-vis chromatogram of mpa-F_430_, two tables related to the protonation of titratable residues in each MCR, and the topologies and associated force field parameters for the F_430_ cofactors, CH_3_-S-CoM, and HS-CoB, are all provided.

## Conflict of Interest Statement

The authors declare no conflict of interest.

## Notes

### Competing Interest Statement

The authors have declared no competing interest.

