## Supporting Information for "Structural dynamics of the methyl-coenzyme M reductase active site are influenced by coenzyme F_430_ modifications"

**Supplemental Methods**

Cultivation of *M. acetivorans*. *Methanosarcina acetivorans* WWM60 was obtained from Dr. Biswarup Mukhopadhyay (Virginia Tech, originally from W. W. Metcalf, University of Illinois at Urbana-Champaign) and was cultured in high-salt medium[1] with acetate (200 mM sodium acetate), methanol (125 mM), or trimethylamine (50 mM) at 37 ºC. A typical $F_{430}$ extraction was performed with cells from 300-500 mL of culture. Bottles with Balch-type closures were used; either 125 mL (Wheaton) bottles containing 75 mL or 1 L (Chemglass) bottles containing 500 mL. The medium was reduced with 0.025% sodium sulfide before inoculation and the headspace contained 80% $N_2$/20% $CO_2$ (10 psi).

Partial purification and analysis of $F_{430}$s. Cells were harvested by centrifugation aerobically and then processed immediately for $F_{430}$ extraction. Pellets (~0.2 g wet weight) were resuspended in 4 mL of water followed by sonication on ice using a Misonex sonicator equipped with a microtip. Two sonication cycles, each one minute, were performed with duty cycle set at 50 (%/1 sec) and the power at 4. Formic acid was added to a final concentration of 1% followed by centrifugation of the acidified lysates at 4,400 x *g*. The supernatant was transferred to a new tube, neutralized with NaOH, and diluted 2x with 50 mM Tris, pH 7.5. The resulting sample was filtered and then applied to a gravity flow column with Q Sepharose Fast Flow resin (2 x 10 cm, Cytiva) equilibrated with 50 mM Tris, pH 7.5. After washing with 10 mL of 50 mM Tris, pH 7.5, the $F_{430}$ was eluted with 10 mL of 20 mM formic acid. The sample was concentrated down to 500 µl under vacuum at 30 ºC, then applied to a 3 kDa MWCO Amicon Ultra concentrator (Millipore-Sigma) to remove any remaining large molecules. The filtrate was further concentrated down to ~100 µL followed by LC-MS or HPLC with diode array (HPLC-DAD) analysis.

For high-resolution LC-MS analysis, a Waters Synapt G2-S HDMS interfaced with an Acquity I-Class UPLC system with an Acquity BEH C18 column (2.1 mm x 50 mm; particle size, 1.7 µm; maintained at 35 ºC) was used. Solvent A was water with 0.1% formic acid, and solvent B was acetonitrile with 0.1% formic acid. The flow rate was 0.2 ml/min, and gradient elution was employed in the following manner (time [min], percent solvent B): (0.01, 1), (5, 20), (7, 95), and (8, 95). Two microliters of sample were injected. The mass spectral data were collected in high-resolution MSe continuum mode (nonselective MS/MS acquisition mode). Parameters were a 2.8-kV capillary voltage, a 125°C source temperature, a 350 °C desolvation temperature, a 35-V sampling cone, 50-liter/h cone gas flow, a 500-liter/h desolvation gas flow, and a 6-liter/h nebulizer gas flow. The collision energies for the low-energy scans (function 1) were 4 V and 2 V in the trap region and the transfer region, respectively. Collision energies for the high-energy scans (function 2) were ramped from 25 to 45 V in the trap region and 2 V in the transfer region. Data were analyzed using MassLynx software (Waters).

For HPLC-DAD analysis, a Shimadzu a Shimadzu HPLC equipped with a photodiode array (PDA) detector and a Kinetex Polar C18 column (Phenomenex, 2.6 µm, 150 x 4.6 mm) was used. The column oven was set at 30°C. Solvent A was 0.1% (v/v) formic acid in water and solvent B was 100% methanol. The flow rate was 0.7 mL $min^{-1}$ and the method consisted of 95% A for 3 min followed by a 20 min linear gradient to 70% B, then a 1 min linear to 100% B followed by 100% B for 3 min. Ten microliters of concentrated cell extract were injected.

Flat-bottom restraints applied during alchemical transformation. In order to ensure a gentle accommodation of the modified $F_{430}$ cofactors by the MCR active-site residues, we employed an alchemical transformation using flat-bottom restraints on key distances and angles that govern the orientation of cofactors within the active site, as shown in Figure S1. Distances were allowed to fluctuate ±1Å from their crystallographic value before the restraint force started to be applied. Two angles were defined to prevent a tilting motion of the $F_{430}$ within the active site: φ was composed by the thioether sulfur atom of $CH_3$-S-CoM, the Ni(I) atom of $F_{430}$ and the pyrrole nitrogen N1 of $F_{430}$ macrocyclic ring, while ψ was composed by the same first two atoms and the pyrrole nitrogen N3 of $F_{430}$. In this way, both angles are almost orthogonal to each other and prevent tilting motions more effectively.

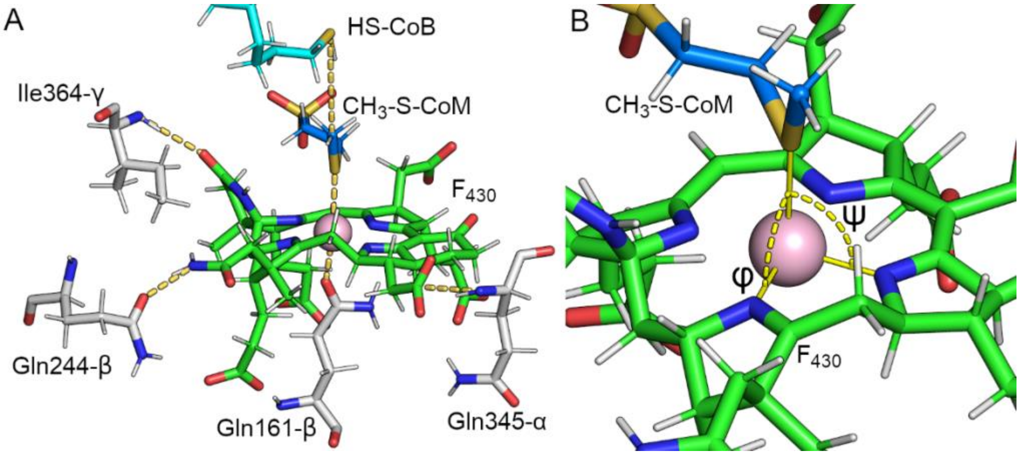

**Figure S1. Flat-bottom restraint scheme applied in this work.** A) Restrained distances between cofactors and protein residues in MCR. B) Restrained angles φ and ψ composed by thioether sulfur, Ni(I) and pyrrole nitrogen N1 and N3, respectively.

<u>Parameters for electric field calculations.</u> TUPÃ uses a configuration file in which is defined the environment set, probe set and other parameters for specific calculation modes. The configuration file used in this work is shown below, where {X} can be A or B, depending on the active site being analyzed. Segment ID COMA and COMB stand for $CH_3$-S-CoM molecules, while protein segment IDs were defined as PROA, PROB, PROC, PROD, PROE and PROF.

```
[Environment Selection]
# The atoms from which we calculate the electric field
sele_environment      = segid PRO* or segid F43{X} or segid COB{X}

[Probe Selection]
# Provide the probe selection for the MODE of your choice
# e.g. if bond is used, then selbond1 and selbond2 must be defined.
mode                 = BOND     # ATOM or BOND or COORDINATE
selbond1             = segid COM{X} and name S2
selbond2             = segid COM{X} and name C3

[Solvent]
include_solvent      = False    # or False
```

```
      [Time]
      dt                  = 10      # Frequency of frames written in your
trajectory (in picosecond)
```

To account only for the contribution of the hydrophobic cage residues to the electric field, we modified the environment set definition to:

```
sele_environment      = (segid PRO{X} and (resid 463 or resid 343 or resid
346)) or (segid PRO{Y} and (resid 359 or resid 365))   #M.acetivorans
```

for *M.acetivorans* MCR, in which {X} and {Y} can be A and C for active site A, or B and D for active site B. For ANME-1 MCR, we used the selection below:

```
sele_environment      = (segid PRO{W} and (resid 462 or resid 344 or resid
347)) or (segid PRO{Z} and (resid 357 or resid 363))   # ANME-1
```

, in which {W} and {Z} can be A and B for active site A, or D and E for active site B.

<u>Force field parameters for MCR cofactors.</u> The parameters below for $CH_3$-S-CoM, HS-CoB and F430 cofactors were obtained from the CGenFF force field version 4.6.

```
* Toppar stream file generated by
* CHARMM General Force Field (CGenFF) program version 2.5
* For use with CGenFF version 4.6
*

read rtf card append
* Topologies generated by
* CHARMM General Force Field (CGenFF) program version 2.5
*
36 1

MASS  -1  Ni1p      58.69300 Ni ! Nickel

! CH3-S-CoM
RESI COM           -1.000 ! param penalty=  21.000 ; charge penalty=   1.050
GROUP             ! CHARGE   CH_PENALTY
ATOM O1     OG2P1  -0.550 !    0.000
ATOM S1     SG3O1   0.736 !    1.050
ATOM O2     OG2P1  -0.550 !    0.000
ATOM O3     OG2P1  -0.550 !    0.000
ATOM C1     CG321  -0.280 !    1.050
ATOM C2     CG321  -0.103 !    1.050
ATOM S2     SG311  -0.113 !    1.050
ATOM H1     HGA2    0.090 !    0.000
ATOM H2     HGA2    0.090 !    0.000
ATOM H3     HGA2    0.090 !    0.000
ATOM H4     HGA2    0.090 !    0.000
ATOM C3     CG331  -0.220 !    0.000
ATOM H5     HGA3    0.090 !    0.000
ATOM H6     HGA3    0.090 !    0.000
ATOM H7     HGA3    0.090 !    0.000

BOND H6   C3
BOND H5   C3
```

```
BOND C3   H7
BOND C3   S2
BOND H1   C1
BOND H2   C1
BOND O3   S1
BOND C1   S1
BOND C1   C2
BOND S2   C2
BOND S1   O1
BOND S1   O2
BOND C2   H3
BOND C2   H4

! HS-CoB
RESI COB         -3.000 ! param penalty=  41.500 ; charge penalty=  37.432
GROUP            ! CHARGE   CH_PENALTY
ATOM O1P    OG2P1  -0.900 !    0.000
ATOM P      PG2     1.101 !    0.474
ATOM O3P    OG2P1  -0.900 !    0.000
ATOM O2P    OG2P1  -0.900 !    0.000
ATOM O4P    OG303  -0.416 !    6.270
ATOM CB     CG311  -0.102 !    8.636
ATOM CG     CG331  -0.281 !    0.379
ATOM CA     CG311   0.019 !   22.180
ATOM C      CG2O3   0.786 !   17.789
ATOM OD1    OG2D2  -0.866 !    1.650
ATOM OD2    OG2D2  -0.866 !    1.650
ATOM N      NG311  -0.692 !   37.432
ATOM C1     CG311   0.290 !   34.068
ATOM O1     OG311  -0.554 !   33.264
ATOM C2     CG321  -0.203 !    6.332
ATOM C3     CG321  -0.191 !    0.330
ATOM C4     CG321  -0.157 !    0.000
ATOM C5     CG321  -0.180 !    0.000
ATOM C6     CG321  -0.174 !    0.000
ATOM C7     CG321  -0.088 !    0.000
ATOM S7     SG311  -0.258 !    0.000
ATOM HB     HGA1    0.090 !    0.060
ATOM HG1    HGA3    0.090 !    0.000
ATOM HG2    HGA3    0.090 !    0.000
ATOM HG3    HGA3    0.090 !    0.000
ATOM HA     HGA1    0.090 !    0.507
ATOM HN     HGPAM1  0.349 !   17.852
ATOM H1     HGA1    0.090 !    0.484
ATOM HO     HGP1    0.403 !   16.270
ATOM H21    HGA2    0.090 !    0.300
ATOM H22    HGA2    0.090 !    0.300
ATOM H31    HGA2    0.090 !    0.000
ATOM H32    HGA2    0.090 !    0.000
ATOM H41    HGA2    0.090 !    0.000
ATOM H42    HGA2    0.090 !    0.000
ATOM H51    HGA2    0.090 !    0.000
ATOM H52    HGA2    0.090 !    0.000
ATOM H61    HGA2    0.090 !    0.000
ATOM H62    HGA2    0.090 !    0.000
ATOM H71    HGA2    0.090 !    0.000
ATOM H72    HGA2    0.090 !    0.000
ATOM HS     HGP3    0.160 !    0.000

BOND O3P  P
BOND O1P  P
BOND P    O2P
BOND P    O4P
BOND O4P  CB
BOND OD1  C
BOND HG1  CG
```

```
BOND HN   N
BOND CB   HB
BOND CB   CG
BOND CB   CA
BOND HG2  CG
BOND C    OD2
BOND C    CA
BOND CG   HG3
BOND N    CA
BOND N    C1
BOND CA   HA
BOND H22  C2
BOND H51  C5
BOND H21  C2
BOND C2   C1
BOND C2   C3
BOND H62  C6
BOND C1   H1
BOND C1   O1
BOND H52  C5
BOND C5   C6
BOND C5   C4
BOND H41  C4
BOND C6   H61
BOND C6   C7
BOND O1   HO
BOND C4   C3
BOND C4   H42
BOND C3   H32
BOND C3   H31
BOND H71  C7
BOND C7   S7
BOND C7   H72
BOND S7   HS
IMPR C      OD1    OD2    CA

RESI F430         -4.000 ! param penalty= 454.500 ; charge penalty= 380.007
GROUP            ! CHARGE   CH_PENALTY
ATOM C1     CG321  -0.186 !   20.762
ATOM C2     CG321  -0.199 !   16.078
ATOM C3     CG2O3   0.620 !    5.293
ATOM C4     CG2O3   0.598 !    8.671
ATOM N1     NG2R50 -0.877 !   33.940
ATOM C5     CG321  -0.259 !    4.975
ATOM C6     CG321  -0.260 !    5.719
ATOM N2     NG2R50 -0.755 !   49.749
ATOM O1     OG2D2  -0.760 !    0.000
ATOM O2     OG2D2  -0.760 !    3.798
ATOM N3     NG2R50 -0.792 !   45.302
ATOM O3     OG2D2  -0.760 !    0.000
ATOM O4     OG2D2  -0.760 !    0.000
ATOM O5     OG2D2  -0.760 !    0.000
ATOM O6     OG2D2  -0.760 !    0.000
ATOM C7     CG321  -0.073 !   25.292
ATOM C8     CG321  -0.206 !   53.058
ATOM C9     CG2DC1  0.465 !  214.872
ATOM C10    CG2DC1  0.431 !  215.580
ATOM C11    CG2R52  0.295 !   39.950
ATOM C12    CG3RC1  0.428 !   31.333
ATOM C13    CG251O  0.151 !  348.225
ATOM C14    CG2R52  0.225 !   56.216
ATOM C15    CG3C50  0.221 !   38.709
ATOM C16    CG3RC1 -0.031 !   26.465
ATOM C17    CG3C51 -0.198 !  238.466
ATOM C18    CG3RC1 -0.074 !   33.642
```

```
ATOM C19    CG3C51 -0.177 !   31.493
ATOM C20    CG3C51 -0.045 !   38.301
ATOM C21    CG3C51 -0.192 !  238.376
ATOM C22    CG3C51 -0.055 !   27.068
ATOM C23    CG3C51  0.204 !   30.794
ATOM C24    CG2R52  0.076 !   55.615
ATOM C25    CG251O  0.213 !  348.253
ATOM C26    CG3C51  0.113 !   30.668
ATOM C27    CG321  -0.150 !   11.702
ATOM C28    CG321  -0.247 !   12.606
ATOM C29    CG321  -0.192 !   24.730
ATOM N4     NG2R53 -0.530 !   38.829
ATOM C30    CG2O1   0.553 !    9.799
ATOM C31    CG2R53  0.288 !   16.765
ATOM C32    CG2O3   0.598 !    8.671
ATOM C33    CG321  -0.143 !   13.703
ATOM C34    CG2O5   0.400 !   25.878
ATOM O7     OG2D1  -0.538 !    4.561
ATOM O8     OG2D1  -0.461 !    0.390
ATOM O9     OG2D2  -0.760 !    3.798
ATOM C35    CG3C52 -0.006 !   12.277
ATOM C36    CG321  -0.187 !   11.511
ATOM N5     NG2S2  -0.621 !    4.561
ATOM O10    OG2D2  -0.760 !    3.798
ATOM O11    OG2D3  -0.411 !   16.265
ATOM C37    CG331  -0.264 !    8.271
ATOM C38    CG331  -0.256 !   15.570
ATOM C39    CG321  -0.260 !    6.152
ATOM C40    CG321  -0.253 !   20.799
ATOM C41    CG2O3   0.620 !    5.293
ATOM C42    CG2O3   0.620 !    5.293
ATOM N6     NG2D1  -0.559 !  380.007
ATOM O12    OG2D2  -0.760 !    0.000
ATOM O13    OG2D2  -0.760 !    3.798
ATOM H1     HGA2    0.090 !    0.040
ATOM H2     HGA2    0.090 !    0.040
ATOM H3     HGA2    0.090 !    0.450
ATOM H4     HGA2    0.090 !    0.450
ATOM H5     HGA2    0.090 !    0.040
ATOM H6     HGA2    0.090 !    0.040
ATOM H7     HGA2    0.090 !    0.040
ATOM H8     HGA2    0.090 !    0.040
ATOM H9     HGA2    0.090 !    2.715
ATOM H10    HGA2    0.090 !    2.715
ATOM H11    HGA2    0.090 !    4.330
ATOM H12    HGA2    0.090 !    4.330
ATOM H13    HGA4    0.122 !   11.791
ATOM H14    HGA1    0.116 !    4.735
ATOM H15    HGA1    0.090 !    2.618
ATOM H16    HGA1    0.090 !    0.906
ATOM H17    HGA1    0.116 !    4.032
ATOM H18    HGA1    0.090 !    2.602
ATOM H19    HGA1    0.090 !    1.030
ATOM H20    HGA1    0.090 !    2.516
ATOM H21    HGA2    0.090 !    0.406
ATOM H22    HGA2    0.090 !    0.406
ATOM H23    HGA2    0.090 !    0.575
ATOM H24    HGA2    0.090 !    0.575
ATOM H25    HGA2    0.090 !    1.200
ATOM H26    HGA2    0.090 !    1.200
ATOM H27    HGP1    0.340 !    2.305
ATOM H28    HGA2    0.090 !    0.450
ATOM H29    HGA2    0.090 !    0.450
ATOM H30    HGA2    0.090 !    0.045
ATOM H31    HGA2    0.090 !    0.045
ATOM H32    HGA2    0.090 !    0.575
```

```
ATOM H33    HGA2    0.090 !    0.575
ATOM H34    HGP1    0.309 !    0.000
ATOM H35    HGP1    0.309 !    0.000
ATOM H36    HGA3    0.090 !    0.415
ATOM H37    HGA3    0.090 !    0.415
ATOM H38    HGA3    0.090 !    0.415
ATOM H39    HGA3    0.090 !    0.020
ATOM H40    HGA3    0.090 !    0.020
ATOM H41    HGA3    0.090 !    0.020
ATOM H42    HGA2    0.090 !    0.040
ATOM H43    HGA2    0.090 !    0.040
ATOM H44    HGA2    0.090 !    0.000
ATOM H45    HGA2    0.090 !    0.000
ATOM H46    HGA1    0.116 !    2.584
ATOM NI     Ni1p    1.000 !

BOND C1   C5
BOND C1   C19
BOND C2   C6
BOND C2   C20
BOND C3   O1
BOND C3   O12
BOND C3   C39
BOND C4   O2
BOND C4   C40
BOND C4   O13
BOND N1   C11
BOND N1   C23
BOND C5   C41
BOND C6   C42
BOND N2   C12
BOND N2   C24
BOND N3   C14
BOND N3   C26
BOND O3   C41
BOND O4   C42
BOND O5   C41
BOND O6   C42
BOND C7   C11
BOND C7   C26
BOND C8   C23
BOND C8   C12
BOND C9   C24
BOND C9   C13
BOND C10  C25
BOND C10  C14
BOND C10  C34
BOND C11  C15
BOND C12  C16
BOND C12  N4
BOND C13  C17
BOND C13  N6
BOND C14  C18
BOND C15  C27
BOND C15  C37
BOND C15  C19
BOND C16  C38
BOND C16  C20
BOND C16  C35
BOND C17  C28
BOND C17  C21
BOND C18  C29
BOND C18  C22
BOND C19  C23
BOND C20  C24
BOND C21  C36
```

```
BOND C21  C25
BOND C22  C26
BOND C22  C40
BOND C25  N6
BOND C27  C30
BOND C28  C32
BOND C29  C33
BOND N4   C31
BOND C30  O7
BOND C30  N5
BOND C31  O8
BOND C31  C35
BOND C32  O9
BOND C32  O10
BOND C33  C34
BOND C34  O11
BOND C36  C39
BOND C1   H1
BOND C1   H2
BOND C2   H3
BOND C2   H4
BOND C5   H5
BOND C5   H6
BOND C6   H7
BOND C6   H8
BOND C7   H9
BOND C7   H10
BOND C8   H11
BOND C8   H12
BOND C9   H13
BOND C17  H14
BOND C18  H15
BOND C19  H16
BOND C21  H17
BOND C22  H18
BOND C23  H19
BOND C26  H20
BOND C27  H21
BOND C27  H22
BOND C28  H23
BOND C28  H24
BOND C29  H25
BOND C29  H26
BOND N4   H27
BOND C33  H28
BOND C33  H29
BOND C35  H30
BOND C35  H31
BOND C36  H32
BOND C36  H33
BOND N5   H34
BOND N5   H35
BOND C37  H36
BOND C37  H37
BOND C37  H38
BOND C38  H39
BOND C38  H40
BOND C38  H41
BOND C39  H42
BOND C39  H43
BOND C40  H44
BOND C40  H45
BOND C20  H46
BOND NI   N1    NI  N2    NI  N3    NI  N6
IMPR C3     O1     O12    C39
IMPR C4     O13    O2     C40
```

```
IMPR C30    C27    N5     O7
IMPR C31    C35    N4     O8
IMPR C32    O10    O9     C28
IMPR C34    C10    C33    O11
IMPR C41    O5     O3     C5
IMPR C42    O6     O4     C6
IMPR N1 C23 C11 NI
IMPR N3 C26 C14 NI
IMPR N2 C24 C12 NI
IMPR N6 C25 C13 NI

! mt-F430
RESI F43T         -5.000 ! param penalty= 454.500 ; charge penalty= 380.007
GROUP            ! CHARGE   CH_PENALTY
ATOM C1     CG321  -0.186 !   20.762
ATOM C2     CG321  -0.199 !   16.078
ATOM C3     CG2O3   0.620 !    5.293
ATOM C4     CG2O3   0.598 !    8.671
ATOM N1     NG2R50 -0.877 !   33.940
ATOM C5     CG321  -0.259 !    4.975
ATOM C6     CG321  -0.260 !    5.719
ATOM N2     NG2R50 -0.755 !   49.749
ATOM O1     OG2D2  -0.760 !    0.000
ATOM O2     OG2D2  -0.760 !    3.798
ATOM N3     NG2R50 -0.792 !   45.302
ATOM O3     OG2D2  -0.760 !    0.000
ATOM O4     OG2D2  -0.760 !    0.000
ATOM O5     OG2D2  -0.760 !    0.000
ATOM O6     OG2D2  -0.760 !    0.000
ATOM C7     CG321  -0.073 !   25.292
ATOM C8     CG321  -0.206 !   53.058
ATOM C9     CG2DC1  0.465 !  214.872
ATOM C10    CG2DC1  0.429 !  215.718
ATOM C11    CG2R52  0.295 !   39.950
ATOM C12    CG3RC1  0.428 !   31.333
ATOM C13    CG251O  0.151 !  348.225
ATOM C14    CG2R52  0.225 !   56.217
ATOM C15    CG3C50  0.221 !   38.709
ATOM C16    CG3RC1 -0.031 !   26.465
ATOM C17    CG3C51 -0.198 !  238.466
ATOM C18    CG3RC1 -0.076 !   33.892
ATOM C19    CG3C51 -0.177 !   31.493
ATOM C20    CG3C51 -0.045 !   38.301
ATOM C21    CG3C51 -0.192 !  238.376
ATOM C22    CG3C51 -0.055 !   27.069
ATOM C23    CG3C51  0.204 !   30.794
ATOM C24    CG2R52  0.076 !   55.615
ATOM C25    CG251O  0.213 !  348.253
ATOM C26    CG3C51  0.113 !   30.668
ATOM C27    CG321  -0.150 !   11.702
ATOM C28    CG321  -0.247 !   12.606
ATOM C29    CG321  -0.029 !   27.064
ATOM N4     NG2R53 -0.530 !   38.829
ATOM C30    CG2O1   0.553 !    9.799
ATOM C31    CG2R53  0.288 !   16.765
ATOM C32    CG2O3   0.598 !    8.671
ATOM C33    CG311   0.109 !   16.585
ATOM C34    CG2O5   0.289 !   27.656
ATOM O7     OG2D1  -0.538 !    4.561
ATOM O8     OG2D1  -0.461 !    0.390
ATOM O9     OG2D2  -0.760 !    3.798
ATOM C35    CG3C52 -0.006 !   12.277
ATOM C36    CG321  -0.187 !   11.511
ATOM N5     NG2S2  -0.621 !    4.561
ATOM O10    OG2D2  -0.760 !    3.798
ATOM O11    OG2D3  -0.370 !   17.050
```

```
ATOM C37    CG331  -0.264 !    8.271
ATOM C38    CG331  -0.256 !   15.570
ATOM C39    CG321  -0.260 !    6.152
ATOM C40    CG321  -0.253 !   20.799
ATOM C41    CG2O3   0.620 !    5.293
ATOM C42    CG2O3   0.620 !    5.293
ATOM N6     NG2D1  -0.559 !  380.007
ATOM O12    OG2D2  -0.760 !    0.000
ATOM O13    OG2D2  -0.760 !    3.798
ATOM H1     HGA2    0.090 !    0.040
ATOM H2     HGA2    0.090 !    0.040
ATOM H3     HGA2    0.090 !    0.450
ATOM H4     HGA2    0.090 !    0.450
ATOM H5     HGA2    0.090 !    0.040
ATOM H6     HGA2    0.090 !    0.040
ATOM H7     HGA2    0.090 !    0.040
ATOM H8     HGA2    0.090 !    0.040
ATOM H9     HGA2    0.090 !    2.715
ATOM H10    HGA2    0.090 !    2.715
ATOM H11    HGA2    0.090 !    4.330
ATOM H12    HGA2    0.090 !    4.330
ATOM H13    HGA4    0.122 !   11.791
ATOM H14    HGA1    0.116 !    4.735
ATOM H15    HGA1    0.090 !    2.618
ATOM H16    HGA1    0.090 !    0.906
ATOM H17    HGA1    0.116 !    4.032
ATOM H18    HGA1    0.090 !    2.602
ATOM H19    HGA1    0.090 !    1.030
ATOM H20    HGA1    0.090 !    2.516
ATOM H21    HGA2    0.090 !    0.406
ATOM H22    HGA2    0.090 !    0.406
ATOM H23    HGA2    0.090 !    0.575
ATOM H24    HGA2    0.090 !    0.575
ATOM H25    HGA2    0.090 !    1.204
ATOM H26    HGA2    0.090 !    1.204
ATOM H27    HGP1    0.340 !    2.305
ATOM H28    HGA1    0.090 !    0.854
ATOM S      SG311  -0.268 !   10.040
ATOM H29    HGA2    0.090 !    0.045
ATOM H30    HGA2    0.090 !    0.045
ATOM H31    HGA2    0.090 !    0.575
ATOM H32    HGA2    0.090 !    0.575
ATOM H33    HGP1    0.309 !    0.000
ATOM H34    HGP1    0.309 !    0.000
ATOM H35    HGA3    0.090 !    0.415
ATOM H36    HGA3    0.090 !    0.415
ATOM H37    HGA3    0.090 !    0.415
ATOM H38    HGA3    0.090 !    0.020
ATOM H39    HGA3    0.090 !    0.020
ATOM H40    HGA3    0.090 !    0.020
ATOM H41    HGA2    0.090 !    0.040
ATOM H42    HGA2    0.090 !    0.040
ATOM H43    HGA2    0.090 !    0.000
ATOM H44    HGA2    0.090 !    0.000
ATOM H45    HGA1    0.116 !    2.584
ATOM C43    CG331  -0.253 !    0.529
ATOM H46    HGA3    0.090 !    0.000
ATOM H47    HGA3    0.090 !    0.000
ATOM H48    HGA3    0.090 !    0.000
ATOM NI     Ni1p    1.000 !

BOND C1   C5
BOND C1   C19
BOND C2   C6
BOND C2   C20
BOND C3   O1
```

```
BOND C3   O12
BOND C3   C39
BOND C4   O2
BOND C4   C40
BOND C4   O13
BOND N1   C11
BOND N1   C23
BOND C5   C41
BOND C6   C42
BOND N2   C12
BOND N2   C24
BOND N3   C14
BOND N3   C26
BOND O3   C41
BOND O4   C42
BOND O5   C41
BOND O6   C42
BOND C7   C11
BOND C7   C26
BOND C8   C23
BOND C8   C12
BOND C9   C24
BOND C9   C13
BOND C10  C25
BOND C10  C14
BOND C10  C34
BOND C11  C15
BOND C12  C16
BOND C12  N4
BOND C13  C17
BOND C13  N6
BOND C14  C18
BOND C15  C27
BOND C15  C37
BOND C15  C19
BOND C16  C38
BOND C16  C20
BOND C16  C35
BOND C17  C28
BOND C17  C21
BOND C18  C29
BOND C18  C22
BOND C19  C23
BOND C20  C24
BOND C21  C36
BOND C21  C25
BOND C22  C26
BOND C22  C40
BOND C25  N6
BOND C27  C30
BOND C28  C32
BOND C29  C33
BOND N4   C31
BOND C30  O7
BOND C30  N5
BOND C31  O8
BOND C31  C35
BOND C32  O9
BOND C32  O10
BOND C33  C34
BOND C34  O11
BOND C36  C39
BOND C1   H1
BOND C1   H2
BOND C2   H3
BOND C2   H4
```

```
BOND C5   H5
BOND C5   H6
BOND C6   H7
BOND C6   H8
BOND C7   H9
BOND C7   H10
BOND C8   H11
BOND C8   H12
BOND C9   H13
BOND C17  H14
BOND C18  H15
BOND C19  H16
BOND C21  H17
BOND C22  H18
BOND C23  H19
BOND C26  H20
BOND C27  H21
BOND C27  H22
BOND C28  H23
BOND C28  H24
BOND C29  H25
BOND C29  H26
BOND N4   H27
BOND C33  H28
BOND C33  S
BOND C35  H29
BOND C35  H30
BOND C36  H31
BOND C36  H32
BOND N5   H33
BOND N5   H34
BOND C37  H35
BOND C37  H36
BOND C37  H37
BOND C38  H38
BOND C38  H39
BOND C38  H40
BOND C39  H41
BOND C39  H42
BOND C40  H43
BOND C40  H44
BOND C20  H45
BOND C43  S
BOND C43  H46
BOND C43  H47
BOND C43  H48
BOND NI   N1    NI  N2    NI  N3    NI  N6
IMPR C3     O1     O12    C39
IMPR C4     O13    O2     C40
IMPR C30    C27    N5     O7
IMPR C31    C35    N4     O8
IMPR C32    O10    O9     C28
IMPR C34    C10    C33    O11
IMPR C41    O5     O3     C5
IMPR C42    O6     O4     C6
IMPR N1 C23 C11 NI
IMPR N3 C26 C14 NI
IMPR N2 C24 C12 NI
IMPR N6 C25 C13 NI

! mpa-F430
RESI F43A          -4.000 ! param penalty= 454.500 ; charge penalty= 380.007
GROUP             ! CHARGE   CH_PENALTY
ATOM C1     CG321  -0.186 !   20.762
```

```
ATOM C2     CG321  -0.199 !   16.078
ATOM C3     CG2O3   0.620 !    5.293
ATOM C4     CG2O3   0.598 !    8.671
ATOM N1     NG2R50 -0.877 !   33.940
ATOM C5     CG321  -0.259 !    4.975
ATOM C6     CG321  -0.260 !    5.719
ATOM N2     NG2R50 -0.755 !   49.749
ATOM O1     OG2D2  -0.760 !    0.000
ATOM O2     OG2D2  -0.760 !    3.798
ATOM N3     NG2R50 -0.792 !   45.302
ATOM O3     OG2D2  -0.760 !    0.000
ATOM O4     OG2D2  -0.760 !    0.000
ATOM O5     OG2D2  -0.760 !    0.000
ATOM O6     OG2D2  -0.760 !    0.000
ATOM C7     CG321  -0.073 !   25.292
ATOM C8     CG321  -0.206 !   53.058
ATOM C9     CG2DC1  0.465 !  214.872
ATOM C10    CG2DC1  0.429 !  215.718
ATOM H1     HGP1    0.309 !    0.000
ATOM C11    CG2R52  0.295 !   39.950
ATOM C12    CG3RC1  0.428 !   31.333
ATOM C13    CG251O  0.151 !  348.225
ATOM C14    CG2R52  0.225 !   56.217
ATOM C15    CG3C50  0.221 !   38.709
ATOM C16    CG3RC1 -0.031 !   26.465
ATOM C17    CG3C51 -0.198 !  238.466
ATOM C18    CG3RC1 -0.076 !   33.892
ATOM C19    CG3C51 -0.177 !   31.493
ATOM C20    CG3C51 -0.045 !   38.301
ATOM C21    CG3C51 -0.192 !  238.376
ATOM C22    CG3C51 -0.055 !   27.069
ATOM C23    CG3C51  0.204 !   30.794
ATOM C24    CG2R52  0.076 !   55.615
ATOM C25    CG251O  0.213 !  348.253
ATOM C26    CG3C51  0.113 !   30.668
ATOM C27    CG321  -0.150 !   11.702
ATOM C28    CG321  -0.247 !   12.606
ATOM C29    CG321  -0.029 !   27.062
ATOM N4     NG2R53 -0.530 !   38.829
ATOM C30    CG2O1   0.553 !    9.799
ATOM C31    CG2R53  0.288 !   16.765
ATOM C32    CG2O3   0.598 !    8.671
ATOM C33    CG311   0.074 !   16.624
ATOM C34    CG2O5   0.289 !   27.673
ATOM O7     OG2D1  -0.538 !    4.561
ATOM O8     OG2D1  -0.461 !    0.390
ATOM O9     OG2D2  -0.760 !    3.798
ATOM C35    CG3C52 -0.006 !   12.277
ATOM C36    CG321  -0.187 !   11.511
ATOM N5     NG2S2  -0.621 !    4.561
ATOM O10    OG2D2  -0.760 !    3.798
ATOM O11    OG2D3  -0.370 !   17.050
ATOM C37    CG331  -0.264 !    8.271
ATOM C38    CG331  -0.256 !   15.570
ATOM C39    CG321  -0.260 !    6.152
ATOM C40    CG321  -0.253 !   20.799
ATOM C41    CG2O3   0.620 !    5.293
ATOM C42    CG2O3   0.620 !    5.293
ATOM N6     NG2D1  -0.559 !  380.007
ATOM O12    OG2D2  -0.760 !    0.000
ATOM O13    OG2D2  -0.760 !    3.798
ATOM H2     HGA2    0.090 !    0.040
ATOM H3     HGA2    0.090 !    0.040
ATOM H4     HGA2    0.090 !    0.450
ATOM H5     HGA2    0.090 !    0.450
ATOM H6     HGA2    0.090 !    0.040
```

```
ATOM H7      HGA2    0.090 !    0.040
ATOM H8      HGA2    0.090 !    0.040
ATOM H9      HGA2    0.090 !    0.040
ATOM H10     HGA2    0.090 !    2.715
ATOM H11     HGA2    0.090 !    2.715
ATOM H12     HGA2    0.090 !    4.330
ATOM H13     HGA2    0.090 !    4.330
ATOM H14     HGA4    0.122 !   11.791
ATOM H15     HGA1    0.116 !    4.735
ATOM H16     HGA1    0.090 !    2.618
ATOM H17     HGA1    0.090 !    0.906
ATOM H18     HGA1    0.116 !    4.032
ATOM H19     HGA1    0.090 !    2.602
ATOM H20     HGA1    0.090 !    1.030
ATOM H21     HGA1    0.090 !    2.516
ATOM H22     HGA2    0.090 !    0.406
ATOM H23     HGA2    0.090 !    0.406
ATOM H24     HGA2    0.090 !    0.575
ATOM H25     HGA2    0.090 !    0.575
ATOM H26     HGA2    0.090 !    1.204
ATOM H27     HGA2    0.090 !    1.204
ATOM H28     HGP1    0.340 !    2.305
ATOM H29     HGA1    0.090 !    0.854
ATOM S       SG311  -0.221 !   10.269
ATOM H30     HGA2    0.090 !    0.045
ATOM H31     HGA2    0.090 !    0.045
ATOM H32     HGA2    0.090 !    0.575
ATOM H33     HGA2    0.090 !    0.575
ATOM H34     HGP1    0.309 !    0.000
ATOM H35     HGP1    0.309 !    0.000
ATOM H36     HGA3    0.090 !    0.415
ATOM H37     HGA3    0.090 !    0.415
ATOM H38     HGA3    0.090 !    0.415
ATOM H39     HGA3    0.090 !    0.020
ATOM H40     HGA3    0.090 !    0.020
ATOM H41     HGA3    0.090 !    0.020
ATOM H42     HGA2    0.090 !    0.040
ATOM H43     HGA2    0.090 !    0.040
ATOM H44     HGA2    0.090 !    0.000
ATOM H45     HGA2    0.090 !    0.000
ATOM H46     HGA1    0.116 !    2.584
ATOM C43     CG321  -0.175 !    2.305
ATOM H47     HGA2    0.090 !    0.000
ATOM H48     HGA2    0.090 !    0.000
ATOM O14     OG2D1  -0.538 !    0.000
ATOM C44     CG321  -0.186 !    2.571
ATOM H49     HGA2    0.090 !    0.000
ATOM H50     HGA2    0.090 !    0.000
ATOM C45     CG2O1   0.547 !    2.391
ATOM N7      NG2S2  -0.621 !    0.000
ATOM H51     HGP1    0.309 !    0.000
ATOM NI      Ni1p    1.000 !

BOND C1   C5
BOND C1   C19
BOND C2   C6
BOND C2   C20
BOND C3   O1
BOND C3   O12
BOND C3   C39
BOND C4   O2
BOND C4   C40
BOND C4   O13
BOND N1   C11
BOND N1   C23
BOND C5   C41
```

```
BOND C6   C42
BOND N2   C12
BOND N2   C24
BOND N3   C14
BOND N3   C26
BOND O3   C41
BOND O4   C42
BOND O5   C41
BOND O6   C42
BOND C7   C11
BOND C7   C26
BOND C8   C23
BOND C8   C12
BOND C9   C24
BOND C9   C13
BOND C10  C25
BOND C10  C14
BOND C10  C34
BOND C11  C15
BOND C12  C16
BOND C12  N4
BOND C13  C17
BOND C13  N6
BOND C14  C18
BOND C15  C27
BOND C15  C37
BOND C15  C19
BOND C16  C38
BOND C16  C20
BOND C16  C35
BOND C17  C28
BOND C17  C21
BOND C18  C29
BOND C18  C22
BOND C19  C23
BOND C20  C24
BOND C21  C36
BOND C21  C25
BOND C22  C26
BOND C22  C40
BOND C25  N6
BOND C27  C30
BOND C28  C32
BOND C29  C33
BOND N4   C31
BOND C30  O7
BOND C30  N5
BOND C31  O8
BOND C31  C35
BOND C32  O9
BOND C32  O10
BOND C33  C34
BOND C34  O11
BOND C36  C39
BOND C44  C45
BOND C45  N7
BOND N7   H51
BOND N7   H1
BOND C1   H2
BOND C1   H3
BOND C2   H4
BOND C2   H5
BOND C5   H6
BOND C5   H7
BOND C6   H8
BOND C6   H9
```

```
BOND C7   H10
BOND C7   H11
BOND C8   H12
BOND C8   H13
BOND C9   H14
BOND C17  H15
BOND C18  H16
BOND C19  H17
BOND C21  H18
BOND C22  H19
BOND C23  H20
BOND C26  H21
BOND C27  H22
BOND C27  H23
BOND C28  H24
BOND C28  H25
BOND C29  H26
BOND C29  H27
BOND N4   H28
BOND C33  H29
BOND C33  S
BOND C35  H30
BOND C35  H31
BOND C36  H32
BOND C36  H33
BOND N5   H34
BOND N5   H35
BOND C37  H36
BOND C37  H37
BOND C37  H38
BOND C38  H39
BOND C38  H40
BOND C38  H41
BOND C39  H42
BOND C39  H43
BOND C40  H44
BOND C40  H45
BOND C20  H46
BOND S    C43
BOND H47  C44
BOND H48  C44
BOND O14  C45
BOND C43  C44
BOND C43  H49
BOND C43  H50
BOND NI   N1    NI  N2    NI  N3    NI  N6
IMPR C3     O1     O12    C39
IMPR C4     O13    O2     C40
IMPR C30    C27    N5     O7
IMPR C31    C35    N4     O8
IMPR C32    O10    O9     C28
IMPR C34    C10    C33    O11
IMPR C41    O5     O3     C5
IMPR C42    O6     O4     C6
IMPR C45    C44    N7     O14
IMPR N1 C23 C11 NI
IMPR N3 C26 C14 NI
IMPR N2 C24 C12 NI
IMPR N6 C25 C13 NI

END

read param card flex append
* Parameters generated by analogy by
* CHARMM General Force Field (CGenFF) program version 2.5
```

```
*

! Penalties lower than 10 indicate the analogy is fair; penalties between 10
! and 50 mean some basic validation is recommended; penalties higher than
! 50 indicate poor analogy and mandate extensive validation/optimization.

ATOMS
MASS  -1  Ni1p      58.69300 Ni ! Nickel

NONBONDED nbxmod  5 atom cdiel fshift vatom vdistance vfswitch -
cutnb 14.0 ctofnb 12.0 ctonnb 10.0 eps 1.0 e14fac 1.0 wmin 1.5

Ni1p 0.0 -0.0200 0.7811

BONDS
! For CoB
CG311  NG311   263.00     1.4740 ! from CG314 NG311, penalty= 1
! For F430 cofactors
CG251O CG3C51  300.00     1.5300 ! from CG2R53 CG3C51, penalty= 64
CG251O NG2D1   500.00     1.3570 ! from CG2DC1 NG2D1, penalty= 225
CG2DC1 CG2R52  345.24     1.4254 ! from CG2DC2 CG2R51, penalty= 5
CG2R52 CG3C50  350.00     1.5100 ! from CG2R51 CG3C50, penalty= 5
CG2R52 CG3C51  350.00     1.5050 ! from CG2R52 CG3C52, penalty= 4
CG2R52 CG3RC1  350.00     1.5050 ! from CG2R52 CG3C52, penalty= 20
CG3C50 CG3C51  195.00     1.5180 ! from CG3C50 CG3C52, penalty= 4
CG3C51 NG2R50  400.00     1.4700 ! from CG3C52 NG2R50, penalty= 4
CG3RC1 NG2R50  400.00     1.4700 ! from CG3C52 NG2R50, penalty= 20
CG2D1  CG3C51  365.00     1.5020 ! from CG2D1 CG321, penalty= 65
CG2DC1 CG3RC1  365.00     1.5020 ! from CG2DC1 CG321, penalty= 75
CG3RC1 OG311   428.00     1.4200 ! from CG3C51 OG311, penalty= 16
CG3RC1 SG311   200.19     1.8095 ! from CG3C51 SG311, penalty= 16
NG2R50 Ni1p    500.00     2.0600 ! HEME - distance from xray
NG2D1  Ni1p    500.00     2.0600 ! HEME - distance from xray

ANGLES
! For CoB
CG2O3  CG311  NG311    43.70    110.00 ! from CG2O2 CG311 NG321, penalty= 7.7
CG311  CG311  NG311    73.00    111.20 ! from CG311 CG311 NG321, penalty= 1.2
CG311  CG311  OG303   115.00    109.70 ! from CG331 CG311 OG303, penalty= 1.5
CG321  CG311  NG311    43.70    112.20 ! from CG321 CG311 NG321, penalty= 1.2
NG311  CG311  OG311    74.45    112.79 ! from OG303 CG311 OG311, penalty= 34.5
NG311  CG311  HGA1     32.40    109.50   50.00   2.13000 ! from NG311 CG314 HGA1, penalty= 1
CG311  NG311  CG311    40.50    109.60 ! from CG321 NG311 CG321, penalty= 1.2
CG311  NG311  HGPAM1   35.00    111.00 ! from CG321 NG311 HGPAM1, penalty= 0.6
! for F430 cofactors
CG2DC1 CG251O CG3C51   40.00    120.40 ! from CG2DC3 CG251O NG2R53, penalty= 271
CG2DC1 CG251O NG2D1    90.00    125.00 ! from CG2DC1 CG2D1O NG2D1, penalty= 285
CG3C51 CG251O NG2D1    40.00    112.00 ! from CG2DC3 CG251O NG2R53, penalty= 309
CG251O CG2DC1 CG2O5    60.00    120.00 ! from CG2DC1 CG2DC1 CG2O5, penalty= 23.5
CG251O CG2DC1 CG2R52  170.00    122.00 ! from CG251O CG2DC1 CG2R51, penalty= 2 ! original value:
k0=29.0
CG2O5  CG2DC1 CG2R52   65.00    123.50 ! from CG2DC2 CG2DC1 CG2O5, penalty= 22.5 ! original
value: k0=48.16
CG2R52 CG2DC1 HGA4     32.00    120.00 ! from CG2R51 CG2DC1 HGA4, penalty= 2
CG2DC1 CG2O5  CG321    35.00    121.00 ! from CG2DC1 CG2O5 CG331, penalty= 0.9
CG2DC1 CG2R52 CG3C51   65.00    123.00 ! from CG321 CG2R51 CG3C52, penalty= 75.4 ! original
value: k0=45.35
CG2DC1 CG2R52 CG3RC1   65.00    119.83 ! from CG321 CG2R51 CG3C52, penalty= 76.1 ! original
value: k0=45.35
CG2DC1 CG2R52 NG2R50   25.00    124.48   19.53   2.51910 ! from CG2DC1 CG2R51 NG2R50, penalty= 5
<<<<<<<<<<<<<<<<<<<<<<<<<<<<<< 25 and 124.5
CG321  CG2R52 CG3C50   50.16    126.5 ! from CG321 CG2R51 CG3C50, penalty= 5
CG3C50 CG2R52 NG2R50  170.00    108.00 ! from CG3C52 CG2R52 NG2R50, penalty= 1.2
CG3C51 CG2R52 NG2R50  170.00    112.00 ! from CG3C52 CG2R52 NG2R50, penalty= 0.4
CG3RC1 CG2R52 NG2R50  170.00    108.00 ! from CG3C52 CG2R52 NG2R50, penalty= 1.1
CG2O1  CG321  CG3C50   52.00    108.00 ! from CG2O1 CG321 CG321, penalty= 10
```

```
CG2O3  CG321  CG3C51   52.00    108.00 ! from CG2O3 CG314 CG3C51, penalty= 5
CG2R52 CG321  CG3C51   58.00    104.00 ! from CG2R52 CG321 CG321, penalty= 10 ! original value:
58.00 and 111.0
CG321  CG321  CG3C51   53.00    112.00    8.00   2.56100 ! from CG321 CG321 CG3C50, penalty= 0.8
CG3C51 CG321  CG3RC1  170.00    111.00    8.00   2.56100 ! from CG321 CG321 CG3RC1, penalty= 10 !
original value: 53.35
CG2R52 CG3C50 CG321    45.00    103.00 ! from CG2R53 CG3C50 CG321, penalty= 1
CG2R52 CG3C50 CG331    45.00    103.00 ! from CG2R53 CG3C50 CG321, penalty= 1.9
CG2R52 CG3C50 CG3C51   70.00    106.50 ! from CG2R53 CG3C51 CG3C52, penalty= 7.4
<<<<<<<<<<<<<<<<<<<<<<<<<<<<<<<<<<<<<<<<<<<<<<
CG321  CG3C50 CG331    58.00    110.00   11.16   2.56100 ! from CG321 CG3C50 CG321, penalty= 0.9
CG321  CG3C50 CG3C51   58.00    115.00    8.00   2.56100 ! from CG321 CG3C51 CG3C51, penalty= 6
CG331  CG3C50 CG3C51   58.00    115.00    8.00   2.56100 ! from CG331 CG3C51 CG3C51, penalty= 6
CG251O CG3C51 CG321    45.00    103.00 ! from CG2R53 CG3C50 CG321, penalty= 9.5
CG251O CG3C51 CG3C51   70.00    106.50 ! from CG2R53 CG3C51 CG3C52, penalty= 3.9
CG251O CG3C51 HGA1     58.00    111.00 ! from CG2R53 CG3C51 HGA1, penalty= 3.5
CG2R52 CG3C51 CG321    45.00    103.00 ! from CG2R53 CG3C50 CG321, penalty= 7
CG2R52 CG3C51 CG3RC1   70.00    106.50 ! from CG2R53 CG3C51 CG3C52, penalty= 2.1
<<<<<<<<<<<<<<<<<<<<<<<<<<<<<<<<<<<<<<<
CG2R52 CG3C51 HGA1     58.00    111.00 ! from CG2R53 CG3C51 HGA1, penalty= 1
CG321  CG3C51 CG3C50   58.00    115.00    8.00   2.56100 ! from CG321 CG3C51 CG3C51, penalty= 0.8
CG321  CG3C51 NG2R50   45.00    112.00 ! from CG2R51 CG3C50 CG331, penalty= 53.4 ! original
value: 103  <<<<
CG3C50 CG3C51 CG3C51   58.00    109.50   11.16   2.56100 ! from CG3C51 CG3C51 CG3C51, penalty=
0.8 <<<<<<<<<<<<<<<<<<<<<<<<<<<<<<<<<<
CG3C50 CG3C51 HGA1     35.00    111.40   22.53   2.17900 ! from CG3C51 CG3C51 HGA1, penalty= 0.8
CG3C51 CG3C51 NG2R50   40.00    107.10 ! from CG3C52 CG3C52 NG2R50, penalty= 4.4
NG2R50 CG3C51 HGA1     44.00    109.80 ! from NG2R50 CG3C52 HGA2, penalty= 4
CG2R53 CG3C52 CG3RC1   70.00    106.50 ! from CG2R53 CG3C52 CG3C52, penalty= 1.1
CG2R52 CG3RC1 CG321    45.00    103.00 ! from CG2R53 CG3C50 CG321, penalty= 23
CG2R52 CG3RC1 CG3C51   70.00    106.50 ! from CG2R53 CG3C51 CG3C52, penalty= 17.4
CG2R52 CG3RC1 HGA1     58.00    111.00 ! from CG2R53 CG3C51 HGA1, penalty= 17
CG321  CG3RC1 NG2R50  120.00    107.00 ! from CG2R51 CG3C50 CG331, penalty= 69.4  ! original
value: 45 and 103
CG321  CG3RC1 NG2R53  120.00    111.00 ! from CG2R51 CG3C50 CG331, penalty= 71.4  ! original
value: 45 and 103
CG331  CG3RC1 CG3C52   58.35    104.00   11.16   2.56100 ! from CG331 CG3RC1 CG3C51, penalty= 0.4
! original value: 113.5
CG3RC1 CG3RC1 NG2R50   70.00    106.00 ! from CG3RC1 CG3RC1 NG2R53, penalty= 19 ! original value:
113.7 <<< N2ring - orig 113.7
NG2R50 CG3RC1 NG2R53   70.00    117.00 ! from NG2R53 CG3RC1 NG2R53, penalty= 19 <<< N4ring - xray
value 116 - original 103
CG251O NG2D1  CG251O  115.00    111.00 ! from CG2DC1 NG2D1 CG2O1, penalty= 454.5 ! original
value: 119.68
CG2R52 NG2R50 CG3C51  115.00    111.00 ! from CG2R52 NG2R50 CG3C52, penalty= 0.4 ! original
value: 102.9
CG2R52 NG2R50 CG3RC1  115.00    111.00 ! from CG2R52 NG2R50 CG3C52, penalty= 1.1 ! original
value: 102.9
CG2D2  CG2D1  CG3C51   48.00    126.00 ! from CG2D2 CG2D1 CG321, penalty= 10
CG3C51 CG2D1  HGA4     40.00    116.00 ! from CG321 CG2D1 HGA4, penalty= 10
CG2D1  CG3C51 CG2R52   68.50    105.00 ! from CG2R51 CG3C51 NG311, penalty= 69
CG2D1  CG3C51 CG3RC1   52.00    112.30 ! from CG2O1 CG3C51 CG3C52, penalty= 30.1
CG2D1  CG3C51 HGA1     50.00    112.00 ! from CG2O1 CG3C51 HGA1, penalty= 29
CG2O5  CG311  CG321    52.00    108.00 ! from CG2O4 CG311 CG321, penalty= 0.5
CG2O5  CG311  SG311    51.61    109.77 ! from CG2O1 CG311 SG311, penalty= 3
CG311  CG321  CG3RC1   53.35    111.00    8.00   2.56100 ! from CG321 CG321 CG3RC1, penalty= 0.6
CG251O CG2DC1 CG3RC1   44.28    118.45 ! from CG251O CG2DC1 CG331, penalty= 14.7
CG2R52 CG2DC1 CG3RC1   48.44    127.34 ! from CG2R53 CG2DC1 CG321, penalty= 14.8
CG3RC1 CG321  CG3RC1   53.35    111.00    8.00   2.56100 ! from CG321 CG321 CG3RC1, penalty= 13.8
CG2O3  CG3C51 CG3RC1   52.00    112.30 ! from CG2O3 CG3C51 CG3C52, penalty= 1.1
CG2DC1 CG3RC1 CG3C51   52.00    112.30 ! from CG2O1 CG3C51 CG3C52, penalty= 45.4
CG2DC1 CG3RC1 CG3RC1   70.00    113.70 ! from CG3RC1 CG3RC1 NG2R61, penalty= 47
CG2DC1 CG3RC1 OG311    45.19    111.93 ! from CG2O1 CG3C50 OG311, penalty= 51
CG321  CG3RC1 SG311    58.00    114.50 ! from CG321 CG321 SG311, penalty= 75
CG3C51 CG3RC1 OG311    75.70    110.10 ! from CG3C51 CG3C51 OG311, penalty= 16
CG3RC1 CG3RC1 OG311    53.35    111.00    8.00   2.56100 ! from CG321 CG3RC1 CG3RC1, penalty= 45
```

```
CG3RC1 CG3RC1 SG311    53.35    111.00    8.00   2.56100 ! from CG321 CG3RC1 CG3RC1, penalty= 113
NG2R50 CG3RC1 NG2R53   70.00    117.00 ! from NG2R53 CG3RC1 NG2R53, penalty= 19
SG311  CG3RC1 HGA1     42.84    107.95 ! from SG311 CG3C51 HGA1, penalty= 16
CG3RC1 OG311  HGP1     50.00    109.00 ! from CG3C51 OG311 HGP1, penalty= 1.5
CG3C52 SG311  CG3RC1   45.20     96.20 ! from CG3C52 SG311 CG3C52, penalty= 1.1
CG2DC1 CG2O5  CG311    35.00    116.00 ! F43A , from CG2DC2 CG2O5 CG331, penalty= 1.5
! HEME dummy
CG251O NG2R50 Ni1p      0.00      0.00 ! HEME
CG3C51 NG2R50 Ni1p      0.00      0.00 ! HEME
CG2R52 NG2R50 Ni1p      0.00      0.00 ! HEME
CG3RC1 NG2R50 Ni1p      0.00      0.00 ! HEME
CG2R52 NG2R50 Ni1p      0.00      0.00 ! HEME
CG321  NG2R50 Ni1p      0.00      0.00 ! HEME
CG251O NG2D1  Ni1p      0.00      0.00 ! HEME
CG3C51 NG2D1  Ni1p      0.00      0.00 ! HEME
CG2R52 NG2D1  Ni1p      0.00      0.00 ! HEME
CG3RC1 NG2D1  Ni1p      0.00      0.00 ! HEME
CG2R52 NG2D1  Ni1p      0.00      0.00 ! HEME
CG321  NG2D1  Ni1p      0.00      0.00 ! HEME
NG2D1  Ni1p   NG2D1     0.00      0.00 ! HEME
NG2R50 Ni1p   NG2R50    0.00      0.00 ! HEME
NG2D1  Ni1p   NG2R50    0.00      0.00 ! HEME

DIHEDRALS
! For CoM
SG311  CG321  CG321  SG3O1      0.1000  3     0.00 ! COM , from SG311 CG321 CG321 SG311, penalty=
21
! For CoB
OG2D2  CG2O3  CG311  NG311      0.5500  2   180.00 ! from OG2D2 CG2O3 CG311 OG301, penalty= 33
CG2O3  CG311  CG311  OG303      2.0000  1   180.00 ! from CG2O2 CG311 CG311 OG301, penalty= 9.5
CG2O3  CG311  CG311  OG303      0.8000  2     0.00 ! from CG2O2 CG311 CG311 OG301, penalty= 9.5
CG331  CG311  CG311  NG311      0.4000  1     0.00 ! from CG331 CG311 CG311 NG321, penalty= 1.2
CG331  CG311  CG311  NG311      0.8000  3     0.00 ! from CG331 CG311 CG311 NG321, penalty= 1.2
NG311  CG311  CG311  OG303      0.4000  1   180.00 ! from NG321 CG311 CG311 OG311, penalty= 16.2
NG311  CG311  CG311  OG303      0.8000  3     0.00 ! from NG321 CG311 CG311 OG311, penalty= 16.2
NG311  CG311  CG311  HGA1       0.5000  3     0.00 ! from NG321 CG311 CG311 HGA1, penalty= 1.2
OG303  CG311  CG311  HGA1       0.1950  3     0.00 ! from OG301 CG311 CG311 HGA1, penalty= 3
NG311  CG311  CG321  CG321      0.1950  3     0.00 ! from NG321 CG311 CG321 CG321, penalty= 1.2
NG311  CG311  CG321  HGA2       0.1600  3     0.00 ! from NG321 CG311 CG321 HGA2, penalty= 1.2
CG2O3  CG311  NG311  CG311      2.0000  1     0.00 ! from CG2O2 CG321 NG311 CG321, penalty= 11.1
CG2O3  CG311  NG311  CG311      1.8000  2     0.00 ! from CG2O2 CG321 NG311 CG321, penalty= 11.1
CG2O3  CG311  NG311  CG311      0.5000  3     0.00 ! from CG2O2 CG321 NG311 CG321, penalty= 11.1
CG2O3  CG311  NG311  HGPAM1     0.6000  1   180.00 ! from CG2O2 CG321 NG311 HGPAM1, penalty= 10.5
CG2O3  CG311  NG311  HGPAM1     0.3000  2     0.00 ! from CG2O2 CG321 NG311 HGPAM1, penalty= 10.5
CG2O3  CG311  NG311  HGPAM1     0.5000  3     0.00 ! from CG2O2 CG321 NG311 HGPAM1, penalty= 10.5
CG311  CG311  NG311  CG311      1.4200  1   180.00 ! from CG321 CG321 NG301 CG321, penalty= 10.2
CG311  CG311  NG311  CG311      0.8200  2     0.00 ! from CG321 CG321 NG301 CG321, penalty= 10.2
CG311  CG311  NG311  CG311      1.0200  3     0.00 ! from CG321 CG321 NG301 CG321, penalty= 10.2
CG311  CG311  NG311  HGPAM1     0.3000  3     0.00 ! from CG321 CG321 NG311 HGPAM1, penalty= 4.6
CG321  CG311  NG311  CG311      1.4200  1   180.00 ! from CG321 CG321 NG301 CG321, penalty= 9.6
CG321  CG311  NG311  CG311      0.8200  2     0.00 ! from CG321 CG321 NG301 CG321, penalty= 9.6
CG321  CG311  NG311  CG311      1.0200  3     0.00 ! from CG321 CG321 NG301 CG321, penalty= 9.6
CG321  CG311  NG311  HGPAM1     0.3000  3     0.00 ! from CG321 CG321 NG311 HGPAM1, penalty= 4
OG311  CG311  NG311  CG311      0.4115  1     0.00 ! from NG3P3 CG314 NG311 CG321, penalty= 41.5
OG311  CG311  NG311  CG311      0.4772  2     0.00 ! from NG3P3 CG314 NG311 CG321, penalty= 41.5
OG311  CG311  NG311  CG311      0.5266  3     0.00 ! from NG3P3 CG314 NG311 CG321, penalty= 41.5
OG311  CG311  NG311  HGPAM1     1.7000  2     0.00 ! from NG311 CG321 NG311 HGPAM1, penalty= 37
OG311  CG311  NG311  HGPAM1     0.3000  3     0.00 ! from NG311 CG321 NG311 HGPAM1, penalty= 37
HGA1   CG311  NG311  CG311      0.0000  3     0.00 ! from HGA1 CG314 NG311 CG321, penalty= 1.6
HGA1   CG311  NG311  HGPAM1     0.0500  3     0.00 ! from HGA1 CG314 NG311 HGPAM1, penalty= 1
CG311  CG311  OG303  PG2        0.4000  1   180.00 ! from CG331 CG311 OG303 PG2, penalty= 1.5
CG311  CG311  OG303  PG2        0.3000  2     0.00 ! from CG331 CG311 OG303 PG2, penalty= 1.5
CG311  CG311  OG303  PG2        0.1000  3     0.00 ! from CG331 CG311 OG303 PG2, penalty= 1.5
NG311  CG311  OG311  HGP1       2.2468  2     0.00 ! from OG303 CG311 OG311 HGP1, penalty= 34.5
NG311  CG311  OG311  HGP1       0.6407  3     0.00 ! from OG303 CG311 OG311 HGP1, penalty= 34.5
! For F430 cofactors
```

```
CG3C51 CG251O CG2DC1 CG2O5      4.6584  2   180.00 ! from NG2R50 CG251O CG2DC1 CG2R61, penalty=
119.5
CG3C51 CG251O CG2DC1 CG2R52     4.5078  2   180.00 ! from NG2R50 CG251O CG2DC1 CG2R51, penalty=
96.5
CG3C51 CG251O CG2DC1 HGA4       5.3882  2   180.00 ! from NG2R50 CG251O CG2DC1 HGA4, penalty=
94.5
NG2D1  CG251O CG2DC1 CG2O5      4.6584  2   180.00 ! from NG2R50 CG251O CG2DC1 CG2R61, penalty=
70
NG2D1  CG251O CG2DC1 CG2R52     4.5078  2   180.00 ! from NG2R50 CG251O CG2DC1 CG2R51, penalty=
47
NG2D1  CG251O CG2DC1 HGA4       5.3882  2   180.00 ! from NG2R50 CG251O CG2DC1 HGA4, penalty= 45
CG2DC1 CG251O CG3C51 CG321      0.0153  3     0.00 ! from CG2R51 CG2R51 CG3C50 CG331, penalty=
143.4
CG2DC1 CG251O CG3C51 CG3C51     0.3500  3   180.00 ! from CG2R51 CG2R51 CG3C51 CG3C51, penalty=
136.5
CG2DC1 CG251O CG3C51 HGA1       0.0000  3     0.00 ! from CG2R51 CG2R51 CG3C51 HGA1, penalty=
136.5
NG2D1  CG251O CG3C51 CG321      1.0000  3   180.00 ! from NG2R53 CG2R53 CG3C50 CG321, penalty=
129
NG2D1  CG251O CG3C51 CG3C51     1.0500  3   180.00 ! from NG2R53 CG2R53 CG3C51 CG3C52, penalty=
123.4
NG2D1  CG251O CG3C51 HGA1       0.0000  3   180.00 ! from NG2R53 CG2R53 CG3C51 HGA1, penalty= 123
CG2DC1 CG251O NG2D1  CG251O     5.5846  2   180.00 ! from CG2DC1 CG2D1O NG2D1 CG2N2, penalty=
359.5
CG3C51 CG251O NG2D1  CG251O     6.6486  1     0.00 ! from CG321 CG2DC1 NG2D1 CG2O1, penalty= 326
CG3C51 CG251O NG2D1  CG251O     5.1386  2   180.00 ! from CG321 CG2DC1 NG2D1 CG2O1, penalty= 326
CG251O CG2DC1 CG2O5  CG321      1.4000  2   180.00 ! from CG2DC3 CG2DC1 CG2O5 CG331, penalty=
26.4
CG251O CG2DC1 CG2O5  OG2D3      1.4000  2   180.00 ! from CG2DC1 CG2DC1 CG2O5 OG2D3, penalty=
23.5
CG2R52 CG2DC1 CG2O5  CG321      1.4000  2   180.00 ! from CG2DC3 CG2DC1 CG2O5 CG331, penalty=
65.4
CG2R52 CG2DC1 CG2O5  OG2D3      1.4000  2   180.00 ! from CG2DC2 CG2DC1 CG2O5 OG2D3, penalty=
22.5
CG251O CG2DC1 CG2R52 CG3C51     0.0503  1   180.00 ! from CG251O CG2DC1 CG2R51 CG2R51, penalty=
100
CG251O CG2DC1 CG2R52 CG3C51     0.7718  2   180.00 ! from CG251O CG2DC1 CG2R51 CG2R51, penalty=
100
CG251O CG2DC1 CG2R52 CG3C51     0.4345  4     0.00 ! from CG251O CG2DC1 CG2R51 CG2R51, penalty=
100
CG251O CG2DC1 CG2R52 CG3RC1     0.0503  1   180.00 ! from CG251O CG2DC1 CG2R51 CG2R51, penalty=
99
CG251O CG2DC1 CG2R52 CG3RC1     0.7718  2   180.00 ! from CG251O CG2DC1 CG2R51 CG2R51, penalty=
99
CG251O CG2DC1 CG2R52 CG3RC1     0.4345  4     0.00 ! from CG251O CG2DC1 CG2R51 CG2R51, penalty=
99
CG251O CG2DC1 CG2R52 NG2R50     1.9911  1   180.00 ! from CG2DC1 CG2DC1 CG2R51 NG2R50, penalty=
28.5
CG251O CG2DC1 CG2R52 NG2R50     1.3259  2   180.00 ! from CG2DC1 CG2DC1 CG2R51 NG2R50, penalty=
28.5
CG251O CG2DC1 CG2R52 NG2R50     0.1046  4     0.00 ! from CG2DC1 CG2DC1 CG2R51 NG2R50, penalty=
28.5
CG2O5  CG2DC1 CG2R52 CG3RC1     1.9911  1   180.00 ! from CG2DC1 CG2DC1 CG2R51 NG2R50, penalty=
167
CG2O5  CG2DC1 CG2R52 CG3RC1     1.3259  2   180.00 ! from CG2DC1 CG2DC1 CG2R51 NG2R50, penalty=
167
CG2O5  CG2DC1 CG2R52 CG3RC1     0.1046  4     0.00 ! from CG2DC1 CG2DC1 CG2R51 NG2R50, penalty=
167
CG2O5  CG2DC1 CG2R52 NG2R50     1.9911  1   180.00 ! from CG2DC1 CG2DC1 CG2R51 NG2R50, penalty=
73.5
CG2O5  CG2DC1 CG2R52 NG2R50     1.3259  2   180.00 ! from CG2DC1 CG2DC1 CG2R51 NG2R50, penalty=
73.5
CG2O5  CG2DC1 CG2R52 NG2R50     0.1046  4     0.00 ! from CG2DC1 CG2DC1 CG2R51 NG2R50, penalty=
73.5
HGA4   CG2DC1 CG2R52 CG3C51     0.0055  2   180.00 ! from HGA4 CG2DC1 CG2R51 NG2R50, penalty=
99.5
```

```
HGA4   CG2DC1 CG2R52 NG2R50     0.0055  2   180.00 ! from HGA4 CG2DC1 CG2R51 NG2R50, penalty= 5
NG2S2  CG2O1  CG321  CG3C50     0.0500  6   180.00 ! from NG2S2 CG2O1 CG321 CG321, penalty= 10
OG2D1  CG2O1  CG321  CG3C50     0.0500  6   180.00 ! from OG2D1 CG2O1 CG321 CG321, penalty= 10
OG2D2  CG2O3  CG321  CG3C51     0.0500  6   180.00 ! from OG2D2 CG2O3 CG314 CG3C51, penalty= 5
CG2DC1 CG2O5  CG321  CG321      0.4000  1     0.00 ! from CG2R61 CG2O5 CG321 CG331, penalty= 21.9
CG2DC1 CG2O5  CG321  CG321      0.1700  2   180.00 ! from CG2R61 CG2O5 CG321 CG331, penalty= 21.9
CG2DC1 CG2O5  CG321  CG321      0.1300  3   180.00 ! from CG2R61 CG2O5 CG321 CG331, penalty= 21.9
CG2DC1 CG2O5  CG321  CG321      0.1000  6   180.00 ! from CG2R61 CG2O5 CG321 CG331, penalty= 21.9
CG2DC1 CG2O5  CG321  HGA2       0.1000  3     0.00 ! from CG2DC1 CG2O5 CG331 HGA3, penalty= 6
CG3C50 CG2R52 CG321  CG3C51     0.1900  3     0.00 ! from NG2R50 CG2R52 CG321 CG321, penalty=
104.5
CG3C50 CG2R52 CG321  HGA2       0.3518  3     0.00 ! from CG3C50 CG2R51 CG321 HGA2, penalty= 5
NG2R50 CG2R52 CG321  CG3C51     0.1900  3     0.00 ! from NG2R50 CG2R52 CG321 CG321, penalty= 10
CG321  CG2R52 CG3C50 CG321      0.3891  3     0.00 ! from CG321 CG2R51 CG3C50 CG331, penalty= 5.9
CG321  CG2R52 CG3C50 CG331      0.3891  3     0.00 ! from CG321 CG2R51 CG3C50 CG331, penalty= 5
CG321  CG2R52 CG3C50 CG3C51     0.3891  3     0.00 ! from CG321 CG2R51 CG3C50 CG331, penalty=
35.9
NG2R50 CG2R52 CG3C50 CG321      2.8000  3   180.00 ! from NG2R50 CG2R52 CG3C52 CG3C52, penalty=
41
NG2R50 CG2R52 CG3C50 CG331      2.8000  3   180.00 ! from NG2R50 CG2R52 CG3C52 CG3C52, penalty=
41
NG2R50 CG2R52 CG3C50 CG3C51     2.8000  3   180.00 ! from NG2R50 CG2R52 CG3C52 CG3C52, penalty=
10.4
CG2DC1 CG2R52 CG3C51 CG321      0.0153  3     0.00 ! from CG2R51 CG2R51 CG3C50 CG331, penalty=
74.4
CG2DC1 CG2R52 CG3C51 CG3RC1     0.3500  3   180.00 ! from CG2R51 CG2R51 CG3C51 CG3C52, penalty=
68.6
CG2DC1 CG2R52 CG3C51 HGA1       0.0000  3     0.00 ! from CG2R51 CG2R51 CG3C51 HGA1, penalty=
67.5
NG2R50 CG2R52 CG3C51 CG321      2.8000  3   180.00 ! from NG2R50 CG2R52 CG3C52 CG3C52, penalty=
35
NG2R50 CG2R52 CG3C51 CG3RC1     2.8000  3   180.00 ! from NG2R50 CG2R52 CG3C52 CG3C52, penalty=
5.1
NG2R50 CG2R52 CG3C51 HGA1       1.4000  3     0.00 ! from NG2R50 CG2R52 CG3C52 HGA2, penalty= 4
CG2DC1 CG2R52 CG3RC1 CG321      0.0153  3     0.00 ! from CG2R51 CG2R51 CG3C50 CG331, penalty=
90.4
CG2DC1 CG2R52 CG3RC1 CG3C51     0.3500  3   180.00 ! from CG2R51 CG2R51 CG3C51 CG3C51, penalty=
83.5
CG2DC1 CG2R52 CG3RC1 HGA1       0.0000  3     0.00 ! from CG2R51 CG2R51 CG3C51 HGA1, penalty=
83.5
NG2R50 CG2R52 CG3RC1 CG321      2.8000  3   180.00 ! from NG2R50 CG2R52 CG3C52 CG3C52, penalty=
51
NG2R50 CG2R52 CG3RC1 CG3C51     2.8000  3   180.00 ! from NG2R50 CG2R52 CG3C52 CG3C52, penalty=
20.4
NG2R50 CG2R52 CG3RC1 HGA1       1.4000  3     0.00 ! from NG2R50 CG2R52 CG3C52 HGA2, penalty= 20
CG2DC1 CG2R52 NG2R50 CG3C51     5.5000  2   180.00 ! from CG2R51 CG2R52 NG2R50 CG3C52, penalty=
62.9
CG2DC1 CG2R52 NG2R50 CG3RC1     5.5000  2   180.00 ! from CG2R51 CG2R52 NG2R50 CG3C52, penalty=
63.6
CG321  CG2R52 NG2R50 CG3C51    17.0000  2   180.00 ! from CG3C52 CG2R52 NG2R50 NG3C51, penalty=
73.5
CG3C50 CG2R52 NG2R50 CG3C51    17.0000  2   180.00 ! from CG3C52 CG2R52 NG2R50 NG3C51, penalty=
43.7
CG3C51 CG2R52 NG2R50 CG3RC1    17.0000  2   180.00 ! from CG3C52 CG2R52 NG2R50 NG3C51, penalty=
41.9
CG3RC1 CG2R52 NG2R50 CG3C51    17.0000  2   180.00 ! from CG3C52 CG2R52 NG2R50 NG3C51, penalty=
43.6
NG2R53 CG2R53 CG3C52 CG3RC1     1.0500  3   180.00 ! from NG2R53 CG2R53 CG3C52 CG3C52, penalty=
1.1
OG2D1  CG2R53 CG3C52 CG3RC1     0.0800  3     0.00 ! from OG2D1 CG2R53 CG3C52 CG3C52, penalty=
1.1
CG3C52 CG2R53 NG2R53 CG3RC1     0.4000  2   180.00 ! from CG3C52 CG2R53 NG2R53 CG3C52, penalty=
1.1
CG2O3  CG321  CG321  CG3C51     0.0645  2     0.00 ! from CG2O3 CG321 CG321 CG321, penalty= 10
CG2O3  CG321  CG321  CG3C51     0.1497  3   180.00 ! from CG2O3 CG321 CG321 CG321, penalty= 10
CG2O3  CG321  CG321  CG3C51     0.0946  4     0.00 ! from CG2O3 CG321 CG321 CG321, penalty= 10
```

```
CG2O3  CG321  CG321  CG3C51     0.1125  5     0.00 ! from CG2O3 CG321 CG321 CG321, penalty= 10
CG2O5  CG321  CG321  CG3RC1     0.2100  1   180.00 ! from CG2O5 CG321 CG321 CG321, penalty= 13.8
CG2O5  CG321  CG321  CG3RC1     0.3900  2     0.00 ! from CG2O5 CG321 CG321 CG321, penalty= 13.8
CG2O5  CG321  CG321  CG3RC1     0.3500  3   180.00 ! from CG2O5 CG321 CG321 CG321, penalty= 13.8
CG2O5  CG321  CG321  CG3RC1     0.1100  4     0.00 ! from CG2O5 CG321 CG321 CG321, penalty= 13.8
CG2O5  CG321  CG321  CG3RC1     0.0900  6   180.00 ! from CG2O5 CG321 CG321 CG321, penalty= 13.8
CG3C51 CG321  CG321  HGA2       0.5000  3     0.00 ! from CG3C50 CG321 CG321 HGA2, penalty= 0.8
CG2O1  CG321  CG3C50 CG2R52     0.8000  4   180.00 ! from CG321 CG321 CG3C50 CG2R53, penalty= 72
CG2O1  CG321  CG3C50 CG331      0.1547  4     0.00 ! from CG2O3 CG314 CG3C51 CG3C52, penalty=
55.5
CG2O1  CG321  CG3C50 CG3C51     0.1547  4     0.00 ! from CG2O3 CG314 CG3C51 CG3C52, penalty=
24.9
HGA2   CG321  CG3C50 CG2R52     0.0000  3     0.00 ! from HGA2 CG321 CG3C50 CG2R53, penalty= 1
HGA2   CG321  CG3C50 CG331      0.0000  3     0.00 ! from HGA2 CG321 CG3C50 CG321, penalty= 0.9
HGA2   CG321  CG3C50 CG3C51     0.1600  3     0.00 ! from HGA2 CG321 CG3C51 CG3C51, penalty= 6
CG2O3  CG321  CG3C51 CG251O     0.1547  4     0.00 ! from CG2O3 CG314 CG3C51 CG3C52, penalty= 80
CG2O3  CG321  CG3C51 CG3C51     0.1547  4     0.00 ! from CG2O3 CG314 CG3C51 CG3C52, penalty= 5.4
CG2O3  CG321  CG3C51 CG3RC1     0.1547  4     0.00 ! from CG2O3 CG314 CG3C51 CG3C52, penalty= 6.1
CG2O3  CG321  CG3C51 HGA1       0.4063  3     0.00 ! from CG2O3 CG314 CG3C51 HGA1, penalty= 5
CG2R52 CG321  CG3C51 CG3C51     0.1547  4     0.00 ! from CG2O3 CG314 CG3C51 CG3C52, penalty=
39.4
CG2R52 CG321  CG3C51 NG2R50     0.2000  3     0.00 ! from NG2S1 CG311 CG321 CG2R51, penalty= 107
CG2R52 CG321  CG3C51 HGA1       0.4063  3     0.00 ! from CG2O3 CG314 CG3C51 HGA1, penalty= 39
CG321  CG321  CG3C51 CG251O     0.8000  4   180.00 ! from CG321 CG321 CG3C50 CG2R53, penalty= 9.5
CG321  CG321  CG3C51 CG2R52     0.8000  4   180.00 ! from CG321 CG321 CG3C50 CG2R53, penalty= 7
CG321  CG321  CG3C51 CG3C50     0.5000  4   180.00 ! from CG321 CG311 CG3C51 CG3C52, penalty= 5.2
CG321  CG321  CG3C51 CG3C51     0.5000  4   180.00 ! from CG321 CG311 CG3C51 CG3C52, penalty= 4.4
CG321  CG321  CG3C51 CG3RC1     0.1500  3     0.00 ! from CG321 CG311 CG3C51 CG3RC1, penalty= 4
CG321  CG321  CG3C51 HGA1       0.1950  3     0.00 ! from CG321 CG311 CG3C51 HGA1, penalty= 4
CG3RC1 CG321  CG3C51 CG3C51     0.5000  4   180.00 ! from CG321 CG311 CG3C51 CG3C52, penalty=
18.2
CG3RC1 CG321  CG3C51 NG2R50     0.8000  4   180.00 ! from CG321 CG321 CG3C50 CG2RC0, penalty=
67.8
CG3RC1 CG321  CG3C51 HGA1       0.1950  3     0.00 ! from CG321 CG311 CG3C51 HGA1, penalty= 17.8
HGA2   CG321  CG3C51 CG251O     0.0000  3     0.00 ! from HGA2 CG321 CG3C50 CG2R53, penalty= 9.5
HGA2   CG321  CG3C51 CG2R52     0.0000  3     0.00 ! from HGA2 CG321 CG3C50 CG2R53, penalty= 7
HGA2   CG321  CG3C51 CG3C50     0.1600  3     0.00 ! from HGA2 CG321 CG3C51 CG3C51, penalty= 0.8
HGA2   CG321  CG3C51 NG2R50     0.0000  3     0.00 ! from HGA2 CG321 CG3C50 CG2RC0, penalty= 54
CG321  CG321  CG3RC1 CG2R52     0.8000  4   180.00 ! from CG321 CG321 CG3C50 CG2R53, penalty= 23
CG3C51 CG321  CG3RC1 CG3RC1     0.1500  3     0.00 ! from CG321 CG321 CG3RC1 CG3RC1, penalty= 10
CG3C51 CG321  CG3RC1 NG2R50     0.8000  4   180.00 ! from CG321 CG321 CG3C50 CG2RC0, penalty= 80
CG3C51 CG321  CG3RC1 NG2R53     0.8000  4   180.00 ! from CG321 CG321 CG3C50 CG2RC0, penalty= 82
HGA2   CG321  CG3RC1 CG2R52     0.0000  3     0.00 ! from HGA2 CG321 CG3C50 CG2R53, penalty= 23
HGA2   CG321  CG3RC1 NG2R50     0.0000  3     0.00 ! from HGA2 CG321 CG3C50 CG2RC0, penalty= 70
HGA2   CG321  CG3RC1 NG2R53     0.0000  3     0.00 ! from HGA2 CG321 CG3C50 CG2RC0, penalty= 72
HGA3   CG331  CG3C50 CG2R52     0.0000  3     0.00 ! from HGA3 CG331 CG3C50 CG2R51, penalty= 2
HGA3   CG331  CG3C50 CG321      0.0000  3     0.00 ! from HGA3 CG331 CG3C50 CG331, penalty= 0.9
HGA3   CG331  CG3C50 CG3C51     0.1600  3     0.00 ! from HGA3 CG331 CG3C51 CG3C51, penalty= 6
HGA3   CG331  CG3RC1 CG3C52     0.1500  3   180.00 ! from HGA3 CG331 CG3RC1 CG3C51, penalty= 0.4
CG2R52 CG3C50 CG3C51 CG321      0.4217  3     0.00 ! from CG331 CG3C51 CG3C52 CG2RC0, penalty=
17.9
CG2R52 CG3C50 CG3C51 CG321      0.5915  4   180.00 ! from CG331 CG3C51 CG3C52 CG2RC0, penalty=
17.9
CG2R52 CG3C50 CG3C51 CG321      0.2301  6   180.00 ! from CG331 CG3C51 CG3C52 CG2RC0, penalty=
17.9
CG2R52 CG3C50 CG3C51 CG3C51     0.3400  3   180.00 ! from CG2R51 CG3C51 CG3C51 CG3C51, penalty= 8
CG2R52 CG3C50 CG3C51 HGA1       0.1900  3     0.00 ! from CG2R51 CG3C51 CG3C51 HGA1, penalty= 8
CG321  CG3C50 CG3C51 CG321      0.0500  3     0.00 ! from CG311 CG3C51 CG3RC1 CG321, penalty=
22.6
CG321  CG3C50 CG3C51 CG3C51     0.1900  3     0.00 ! from CG321 CG3C51 CG3C51 CG3C51, penalty= 6
CG321  CG3C50 CG3C51 HGA1       0.1900  3     0.00 ! from CG321 CG3C51 CG3C51 HGA1, penalty= 6
CG331  CG3C50 CG3C51 CG321      0.2000  3     0.00 ! from CG331 CG3C51 CG3RC1 CG321, penalty= 22
CG331  CG3C50 CG3C51 CG3C51     0.1900  3     0.00 ! from CG331 CG3C51 CG3C51 CG3C51, penalty= 6
CG331  CG3C50 CG3C51 HGA1       0.1900  3     0.00 ! from CG331 CG3C51 CG3C51 HGA1, penalty= 6
CG251O CG3C51 CG3C51 CG251O     0.0075  3     0.00 ! from CG2R53 CG3C51 CG3C52 CG2R53, penalty=
11
```

```
CG251O CG3C51 CG3C51 CG321      0.4217  3     0.00 ! from CG331 CG3C51 CG3C52 CG2RC0, penalty=
12.9
CG251O CG3C51 CG3C51 CG321      0.5915  4   180.00 ! from CG331 CG3C51 CG3C52 CG2RC0, penalty=
12.9
CG251O CG3C51 CG3C51 CG321      0.2301  6   180.00 ! from CG331 CG3C51 CG3C52 CG2RC0, penalty=
12.9
CG251O CG3C51 CG3C51 HGA1       0.1900  3     0.00 ! from CG2R51 CG3C51 CG3C51 HGA1, penalty= 4
CG321  CG3C51 CG3C51 CG321      0.0500  3     0.00 ! from CG311 CG3C51 CG3RC1 CG321, penalty=
16.6
CG321  CG3C51 CG3C51 CG3C50     0.1900  3     0.00 ! from CG321 CG3C51 CG3C51 CG3C51, penalty=
0.8
CG321  CG3C51 CG3C51 CG3RC1     0.1900  3     0.00 ! from CG321 CG3C51 CG3C51 CG3C52, penalty=
1.1
CG321  CG3C51 CG3C51 NG2R50     0.5000  2   180.00 ! from CG331 CG3C51 CG3C52 NG2S0, penalty=
42.9
CG3C50 CG3C51 CG3C51 NG2R50     0.0000  3     0.00 ! from CG3C51 CG3C51 CG3C51 NG2R51, penalty=
25.8
CG3C50 CG3C51 CG3C51 HGA1       0.1900  3     0.00 ! from CG3C51 CG3C51 CG3C51 HGA1, penalty= 0.8
CG3RC1 CG3C51 CG3C51 NG2R50     0.0000  3     0.00 ! from CG3C52 CG3C51 CG3C51 NG2R51, penalty=
26.1
NG2R50 CG3C51 CG3C51 HGA1       0.3000  3     0.00 ! from NG2R50 CG3C52 CG3C52 HGA2, penalty= 8
CG2R52 CG3C51 CG3RC1 CG331      0.4217  3     0.00 ! from CG331 CG3C51 CG3C52 CG2RC0, penalty= 27
CG2R52 CG3C51 CG3RC1 CG331      0.5915  4   180.00 ! from CG331 CG3C51 CG3C52 CG2RC0, penalty= 27
CG2R52 CG3C51 CG3RC1 CG331      0.2301  6   180.00 ! from CG331 CG3C51 CG3C52 CG2RC0, penalty= 27
CG2R52 CG3C51 CG3RC1 CG3C52     0.3400  3   180.00 ! from CG2R51 CG3C51 CG3C51 CG3C51, penalty=
18.4
CG2R52 CG3C51 CG3RC1 CG3RC1     0.1500  3     0.00 ! from NG2R51 CG3C51 CG3RC1 CG3RC1, penalty=
52
CG321  CG3C51 CG3RC1 CG2R52     0.4217  3     0.00 ! from CG331 CG3C51 CG3C52 CG2RC0, penalty=
27.9
CG321  CG3C51 CG3RC1 CG2R52     0.5915  4   180.00 ! from CG331 CG3C51 CG3C52 CG2RC0, penalty=
27.9
CG321  CG3C51 CG3RC1 CG2R52     0.2301  6   180.00 ! from CG331 CG3C51 CG3C52 CG2RC0, penalty=
27.9
CG321  CG3C51 CG3RC1 CG321      0.0500  3     0.00 ! from CG311 CG3C51 CG3RC1 CG321, penalty= 0.6
CG321  CG3C51 CG3RC1 CG331      0.1580  3     0.00 ! from CG311 CG3C51 CG3RC1 CG331, penalty= 0.6
CG321  CG3C51 CG3RC1 CG3C52     0.1900  3     0.00 ! from CG321 CG3C51 CG3C51 CG3C52, penalty= 16
CG3C51 CG3C51 CG3RC1 CG2R52     0.3400  3   180.00 ! from CG2R51 CG3C51 CG3C51 CG3C51, penalty=
18
CG3C51 CG3C51 CG3RC1 CG321      2.2000  2   180.00 ! from CG3C52 CG3C51 CG3RC1 CG321, penalty=
0.4
CG3C51 CG3C51 CG3RC1 CG321      4.0000  3     0.00 ! from CG3C52 CG3C51 CG3RC1 CG321, penalty=
0.4
CG3C51 CG3C51 CG3RC1 CG321      0.5500  6   180.00 ! from CG3C52 CG3C51 CG3RC1 CG321, penalty=
0.4
HGA1   CG3C51 CG3RC1 CG2R52     0.1900  3     0.00 ! from CG2R51 CG3C51 CG3C51 HGA1, penalty= 18
CG321  CG3C51 NG2R50 CG2R52     1.9000  3   180.00 ! from CG3C52 CG3C52 NG2R50 CG2R53, penalty=
36
CG3C51 CG3C51 NG2R50 CG2R52     1.9000  3   180.00 ! from CG3C52 CG3C52 NG2R50 CG2R53, penalty=
5.4
HGA1   CG3C51 NG2R50 CG2R52     0.0000  3     0.00 ! from HGA2 CG3C52 NG2R50 CG2R52, penalty= 4
CG2R53 CG3C52 CG3RC1 CG331      0.4217  3     0.00 ! from CG331 CG3C51 CG3C52 CG2RC0, penalty= 23
CG2R53 CG3C52 CG3RC1 CG331      0.5915  4   180.00 ! from CG331 CG3C51 CG3C52 CG2RC0, penalty= 23
CG2R53 CG3C52 CG3RC1 CG331      0.2301  6   180.00 ! from CG331 CG3C51 CG3C52 CG2RC0, penalty= 23
CG2R53 CG3C52 CG3RC1 CG3C51     0.3400  3   180.00 ! from CG3C52 CG3C51 CG3C52 CG2R51, penalty=
18.9
CG2R53 CG3C52 CG3RC1 CG3RC1     0.1500  3     0.00 ! from NG2R51 CG3C51 CG3RC1 CG3RC1, penalty=
56
HGA2   CG3C52 CG3RC1 CG331      0.1950  1     0.00 ! from HGA2 CG3C52 CG3RC1 CG321, penalty= 0.9
CG331  CG3RC1 CG3RC1 NG2R50     0.1500  3     0.00 ! from CG3C52 CG3RC1 CG3RC1 NG2R51, penalty=
56
CG331  CG3RC1 CG3RC1 NG2R53     0.1500  3     0.00 ! from CG3C52 CG3RC1 CG3RC1 NG2R51, penalty=
40
CG3C51 CG3RC1 CG3RC1 NG2R50     0.1500  3     0.00 ! from CG3C52 CG3RC1 CG3RC1 NG2R51, penalty=
25.4
```

```
CG3C51 CG3RC1 CG3RC1 NG2R53     0.1500  3     0.00 ! from CG3C52 CG3RC1 CG3RC1 NG2R51, penalty=
9.4
CG3C52 CG3RC1 CG3RC1 NG2R50     0.1500  3     0.00 ! from CG3C52 CG3RC1 CG3RC1 NG2R51, penalty=
25
CG3C52 CG3RC1 CG3RC1 NG2R53     0.1500  3     0.00 ! from CG3C52 CG3RC1 CG3RC1 NG2R51, penalty= 9
CG321  CG3RC1 NG2R50 CG2R52     1.9000  3   180.00 ! from CG3C52 CG3C52 NG2R50 CG2R53, penalty=
52
CG3RC1 CG3RC1 NG2R50 CG2R52     1.9000  3   180.00 ! from CG3C52 CG3C52 NG2R50 CG2R53, penalty=
62.1
NG2R53 CG3RC1 NG2R50 CG2R52     2.0000  2   180.00 ! from CG2R51 CG3C52 NG2R50 CG2R52, penalty=
68.5
CG321  CG3RC1 NG2R53 CG2R53     2.3100  3   180.00 ! from CG3C52 CG3C52 NG2R53 CG2R53, penalty=
51
CG321  CG3RC1 NG2R53 HGP1       0.7600  3     0.00 ! from CG3C52 CG3C52 NG2R53 HGP1, penalty= 51
NG2R50 CG3RC1 NG2R53 CG2R53     2.3100  3   180.00 ! from NG2R53 CG3RC1 NG2R53 CG2R53, penalty=
19
NG2R50 CG3RC1 NG2R53 HGP1       0.7600  3     0.00 ! from NG2R53 CG3RC1 NG2R53 HGP1, penalty= 19
CG3C51 CG2D1  CG2D2  HGA5       5.2000  2   180.00 ! from CG321 CG2D1 CG2D2 HGA5, penalty= 10
CG2D2  CG2D1  CG3C51 CG2R52     1.2000  1   180.00 ! from CG2D2 CG2D1 CG321 CG2D1, penalty= 107
CG2D2  CG2D1  CG3C51 CG2R52     0.4000  2   180.00 ! from CG2D2 CG2D1 CG321 CG2D1, penalty= 107
CG2D2  CG2D1  CG3C51 CG2R52     1.3000  3   180.00 ! from CG2D2 CG2D1 CG321 CG2D1, penalty= 107
CG2D2  CG2D1  CG3C51 CG3RC1     0.5000  1   180.00 ! from CG2D2 CG2D1 CG321 CG321, penalty= 98.8
CG2D2  CG2D1  CG3C51 CG3RC1     1.3000  3   180.00 ! from CG2D2 CG2D1 CG321 CG321, penalty= 98.8
CG2D2  CG2D1  CG3C51 HGA1       0.1200  3     0.00 ! from CG2D2 CG2D1 CG321 HGA2, penalty= 65
HGA4   CG2D1  CG3C51 CG2R52     0.0000  3     0.00 ! from HGA4 CG2D1 CG321 CG2D1, penalty= 107
HGA4   CG2D1  CG3C51 CG3RC1     0.1200  3     0.00 ! from HGA4 CG2D1 CG321 CG321, penalty= 98.8
HGA4   CG2D1  CG3C51 HGA1       0.0000  3     0.00 ! from HGA4 CG2D1 CG321 HGA2, penalty= 65
CG2DC1 CG2R52 CG3C51 CG2D1      1.7982  3   180.00 ! from CG2R51 CG2R51 CG3C51 CG2O1, penalty=
96.5
NG2R50 CG2R52 CG3C51 CG2D1      3.5000  3   180.00 ! from NG2R50 CG2R52 CG3C52 CG2R51, penalty=
46.5
CG2D1  CG3C51 CG3RC1 CG331      0.4217  3     0.00 ! from CG331 CG3C51 CG3C52 CG2RC0, penalty= 64
CG2D1  CG3C51 CG3RC1 CG331      0.5915  4   180.00 ! from CG331 CG3C51 CG3C52 CG2RC0, penalty= 64
CG2D1  CG3C51 CG3RC1 CG331      0.2301  6   180.00 ! from CG331 CG3C51 CG3C52 CG2RC0, penalty= 64
CG2D1  CG3C51 CG3RC1 CG3C52     0.1400  3     0.00 ! from CG2O1 CG3C51 CG3C52 CG3C52, penalty= 49
CG2D1  CG3C51 CG3RC1 CG3RC1     0.1500  3     0.00 ! from NG2R61 CG3C51 CG3RC1 CG3RC1, penalty=
47
CG251O CG2DC1 CG2O5  CG311      1.4000  2   180.00 ! from CG2DC3 CG2DC1 CG2O5 CG331, penalty= 27
CG2R52 CG2DC1 CG2O5  CG311      1.4000  2   180.00 ! from CG2DC3 CG2DC1 CG2O5 CG331, penalty= 66
CG2DC1 CG2O5  CG311  CG321      0.4000  1     0.00 ! from CG2R61 CG2O5 CG321 CG331, penalty= 25.9
CG2DC1 CG2O5  CG311  CG321      0.1700  2   180.00 ! from CG2R61 CG2O5 CG321 CG331, penalty= 25.9
CG2DC1 CG2O5  CG311  CG321      0.1300  3   180.00 ! from CG2R61 CG2O5 CG321 CG331, penalty= 25.9
CG2DC1 CG2O5  CG311  CG321      0.1000  6   180.00 ! from CG2R61 CG2O5 CG321 CG331, penalty= 25.9
CG2DC1 CG2O5  CG311  SG311      0.0082  2   180.00 ! from OG2D3 CG2O5 CG321 SG311, penalty= 59.5
CG2DC1 CG2O5  CG311  SG311      0.1968  3   180.00 ! from OG2D3 CG2O5 CG321 SG311, penalty= 59.5
CG2DC1 CG2O5  CG311  HGA1       0.1000  3     0.00 ! from CG2DC1 CG2O5 CG331 HGA3, penalty= 11
OG2D3  CG2O5  CG311  CG321      0.7500  1   180.00 ! from OG2D3 CG2O5 CG321 CG321, penalty= 4
OG2D3  CG2O5  CG311  CG321      0.1800  2   180.00 ! from OG2D3 CG2O5 CG321 CG321, penalty= 4
OG2D3  CG2O5  CG311  CG321      0.0650  3   180.00 ! from OG2D3 CG2O5 CG321 CG321, penalty= 4
OG2D3  CG2O5  CG311  CG321      0.0300  6     0.00 ! from OG2D3 CG2O5 CG321 CG321, penalty= 4
OG2D3  CG2O5  CG311  SG311      0.0082  2   180.00 ! from OG2D3 CG2O5 CG321 SG311, penalty= 4
OG2D3  CG2O5  CG311  SG311      0.1968  3   180.00 ! from OG2D3 CG2O5 CG321 SG311, penalty= 4
CG2O5  CG311  CG321  CG3RC1     0.2000  3     0.00 ! from CG2O2 CG311 CG321 CG321, penalty= 15.8
CG2O5  CG311  CG321  HGA2       0.2000  3     0.00 ! from CG2O4 CG311 CG321 HGA2, penalty= 0.5
SG311  CG311  CG321  CG3RC1     0.1950  3     0.00 ! from SG311 CG311 CG321 CG321, penalty= 13.8
HGA1   CG311  CG321  CG3RC1     0.1500  3     0.00 ! from CG3RC1 CG311 CG311 HGA1, penalty= 4
CG2O5  CG311  SG311  CG331      1.5818  1   180.00 ! from CG2O1 CG311 SG311 CG331, penalty= 3
CG2O5  CG311  SG311  CG331      0.6801  3     0.00 ! from CG2O1 CG311 SG311 CG331, penalty= 3
CG321  CG311  SG311  CG331      0.2400  1   180.00 ! from CG321 CG311 SG311 CG321, penalty= 0.9
CG321  CG311  SG311  CG331      0.3700  3     0.00 ! from CG321 CG311 SG311 CG321, penalty= 0.9
CG311  CG321  CG3RC1 CG2R52     0.8000  4   180.00 ! from CG321 CG321 CG3C50 CG2R53, penalty=
23.6
CG311  CG321  CG3RC1 CG3C51     0.1580  3     0.00 ! from CG321 CG321 CG3RC1 CG3C51, penalty= 0.6
CG311  CG321  CG3RC1 HGA1       0.1500  3     0.00 ! from CG321 CG321 CG3RC1 HGA1, penalty= 0.6
CG3C51 CG251O CG2DC1 CG3RC1     4.6584  2   180.00 ! from NG2R50 CG251O CG2DC1 CG331, penalty=
109.2
```

```
NG2D1  CG251O CG2DC1 CG3RC1     4.6584  2   180.00 ! from NG2R50 CG251O CG2DC1 CG331, penalty=
59.7
CG3RC1 CG2DC1 CG2R52 CG3RC1     0.9539  2   180.00 ! from CG321 CG2DC1 CG2R53 NG2R51, penalty=
129.3
CG3RC1 CG2DC1 CG2R52 CG3RC1     0.6267  4     0.00 ! from CG321 CG2DC1 CG2R53 NG2R51, penalty=
129.3
CG3RC1 CG2DC1 CG2R52 NG2R50     0.5016  2   180.00 ! from CG321 CG2DC1 CG2R53 NG2R50, penalty=
36.8
CG3RC1 CG2DC1 CG2R52 NG2R50     0.4507  3   180.00 ! from CG321 CG2DC1 CG2R53 NG2R50, penalty=
36.8
CG3RC1 CG2DC1 CG2R52 NG2R50     0.0954  6     0.00 ! from CG321 CG2DC1 CG2R53 NG2R50, penalty=
36.8
CG251O CG2DC1 CG3RC1 CG3C51     0.5000  2     0.00 ! from CG2DC1 CG2DC1 CG321 CG321, penalty=
128.5
CG251O CG2DC1 CG3RC1 CG3C51     0.3000  3     0.00 ! from CG2DC1 CG2DC1 CG321 CG321, penalty=
128.5
CG251O CG2DC1 CG3RC1 CG3RC1     0.5000  2     0.00 ! from CG2DC1 CG2DC1 CG321 CG321, penalty=
172.3
CG251O CG2DC1 CG3RC1 CG3RC1     0.3000  3     0.00 ! from CG2DC1 CG2DC1 CG321 CG321, penalty=
172.3
CG251O CG2DC1 CG3RC1 OG311      1.9000  1   180.00 ! from CG2DC1 CG2DC1 CG321 OG311, penalty=
98.5
CG251O CG2DC1 CG3RC1 OG311      0.4000  2   180.00 ! from CG2DC1 CG2DC1 CG321 OG311, penalty=
98.5
CG251O CG2DC1 CG3RC1 OG311      0.6000  3   180.00 ! from CG2DC1 CG2DC1 CG321 OG311, penalty=
98.5
CG2R52 CG2DC1 CG3RC1 CG3C51     0.1900  3     0.00 ! from CG2R53 CG2DC1 CG321 CG321, penalty= 106
CG2R52 CG2DC1 CG3RC1 CG3RC1     0.1900  3     0.00 ! from CG2R53 CG2DC1 CG321 CG321, penalty=
149.8
CG2R52 CG2DC1 CG3RC1 OG311      0.1900  3     0.00 ! from CG2R53 CG2DC1 CG321 CG321, penalty= 121
OG2D2  CG2O3  CG3C51 CG3RC1     0.1600  3     0.00 ! from OG2D2 CG2O3 CG3C51 CG3C52, penalty= 1.1
CG3RC1 CG321  CG3RC1 CG2R52     0.8000  4   180.00 ! from CG321 CG321 CG3C50 CG2R53, penalty=
36.8
CG3RC1 CG321  CG3RC1 CG3C51     0.1580  3     0.00 ! from CG321 CG321 CG3RC1 CG3C51, penalty=
13.8
CG3RC1 CG321  CG3RC1 CG3RC1     0.1500  3     0.00 ! from CG321 CG321 CG3RC1 CG3RC1, penalty=
13.8
CG3RC1 CG321  CG3RC1 SG311      0.1950  3     0.00 ! from CG321 CG321 CG321 SG311, penalty= 88.8
CG3RC1 CG321  CG3RC1 HGA1       0.1500  3     0.00 ! from CG321 CG321 CG3RC1 HGA1, penalty= 13.8
HGA2   CG321  CG3RC1 SG311      0.0100  3     0.00 ! from SG311 CG321 CG321 HGA2, penalty= 75
CG2O3  CG3C51 CG3C52 SG311      0.1403  3     0.00 ! from CG2O1 CG3C51 CG3C52 SG311, penalty= 7.5
CG3RC1 CG3C51 CG3C52 SG311      0.0891  3     0.00 ! from CG3C51 CG3C51 CG3C51 SG311, penalty=
5.5
CG2O3  CG3C51 CG3RC1 CG2DC1     3.4478  3     0.00 ! from NG2S1 CG3C51 CG3C52 CG2R53, penalty=
112.5
CG2O3  CG3C51 CG3RC1 CG3RC1     0.1500  3     0.00 ! from NG2R61 CG3C51 CG3RC1 CG3RC1, penalty=
52
CG2O3  CG3C51 CG3RC1 OG311      0.1400  3     0.00 ! from CG2R51 CG3C51 CG3C51 OG311, penalty=
66.5
CG3C52 CG3C51 CG3RC1 CG2DC1     0.1400  3     0.00 ! from CG2O1 CG3C51 CG3C52 CG3C52, penalty= 49
CG3C52 CG3C51 CG3RC1 OG311      0.2000  3     0.00 ! from CG3C52 CG3C51 CG3C51 OG311, penalty= 16
HGA1   CG3C51 CG3RC1 CG2DC1     0.1400  3     0.00 ! from CG2O1 CG3C51 CG3C52 HGA2, penalty= 49
HGA1   CG3C51 CG3RC1 OG311      0.1950  3     0.00 ! from OG311 CG3C51 CG3C51 HGA1, penalty= 16
CG3C51 CG3C52 SG311  CG3RC1     0.1718  1   180.00 ! from CG3C51 CG3C52 SG311 CG3C52, penalty=
1.1
CG3C51 CG3C52 SG311  CG3RC1     0.3700  3     0.00 ! from CG3C51 CG3C52 SG311 CG3C52, penalty=
1.1
HGA2   CG3C52 SG311  CG3RC1     0.2771  3     0.00 ! from HGA2 CG3C52 SG311 CG3C52, penalty= 1.1
CG2DC1 CG3RC1 CG3RC1 CG321      0.1500  3     0.00 ! from CG321 CG3RC1 CG3RC1 CG321, penalty= 70
CG2DC1 CG3RC1 CG3RC1 SG311      0.1500  3     0.00 ! from CG321 CG3RC1 CG3RC1 CG321, penalty= 183
CG2DC1 CG3RC1 CG3RC1 HGA1       0.1500  3     0.00 ! from NG2R61 CG3RC1 CG3RC1 HGA1, penalty= 47
CG321  CG3RC1 CG3RC1 OG311      0.1500  3     0.00 ! from CG321 CG3RC1 CG3RC1 CG321, penalty= 45
CG3C51 CG3RC1 CG3RC1 SG311      0.0500  3     0.00 ! from CG321 CG3RC1 CG3RC1 CG3C51, penalty=
113
OG311  CG3RC1 CG3RC1 SG311      0.1500  3     0.00 ! from CG321 CG3RC1 CG3RC1 CG321, penalty= 158
OG311  CG3RC1 CG3RC1 HGA1       0.3000  3     0.00 ! from OG3C31 CG3RC1 CG3RC1 HGA1, penalty= 31
```

```
CG2DC1 CG3RC1 OG311  HGP1       0.0292  1     0.00 ! from CG2O1 CG3C50 OG311 HGP1, penalty= 51
CG2DC1 CG3RC1 OG311  HGP1       1.3663  2     0.00 ! from CG2O1 CG3C50 OG311 HGP1, penalty= 51
CG2DC1 CG3RC1 OG311  HGP1       0.5143  3     0.00 ! from CG2O1 CG3C50 OG311 HGP1, penalty= 51
CG3C51 CG3RC1 OG311  HGP1       0.2900  1     0.00 ! from CG3C51 CG3C51 OG311 HGP1, penalty= 16
CG3C51 CG3RC1 OG311  HGP1       0.6200  2     0.00 ! from CG3C51 CG3C51 OG311 HGP1, penalty= 16
CG3C51 CG3RC1 OG311  HGP1       0.0500  3     0.00 ! from CG3C51 CG3C51 OG311 HGP1, penalty= 16
CG3RC1 CG3RC1 OG311  HGP1       1.5000  1     0.00 ! from CG3RC1 CG3C51 OG311 HGP1, penalty= 56
CG3RC1 CG3RC1 OG311  HGP1       0.3000  2   180.00 ! from CG3RC1 CG3C51 OG311 HGP1, penalty= 56
CG3RC1 CG3RC1 OG311  HGP1       0.3200  3     0.00 ! from CG3RC1 CG3C51 OG311 HGP1, penalty= 56
CG321  CG3RC1 SG311  CG3C52     0.1718  1   180.00 ! from CG3C51 CG3C52 SG311 CG3C52, penalty=
51.4
CG321  CG3RC1 SG311  CG3C52     0.3700  3     0.00 ! from CG3C51 CG3C52 SG311 CG3C52, penalty=
51.4
CG3RC1 CG3RC1 SG311  CG3C52     0.1718  1   180.00 ! from CG3C51 CG3C52 SG311 CG3C52, penalty=
61.5
CG3RC1 CG3RC1 SG311  CG3C52     0.3700  3     0.00 ! from CG3C51 CG3C52 SG311 CG3C52, penalty=
61.5
HGA1   CG3RC1 SG311  CG3C52     0.2771  3     0.00 ! from HGA2 CG3C52 SG311 CG3C52, penalty= 20
CG2O5  CG311  SG311  CG321      1.5818  1   180.00 ! F43A , from CG2O1 CG311 SG311 CG321,
penalty= 3
CG2O5  CG311  SG311  CG321      0.6801  3     0.00 ! F43A , from CG2O1 CG311 SG311 CG321,
penalty= 3
CG2O1  CG321  CG321  SG311      0.2000  3     0.00 ! F43A , from CG2O1 CG311 CG321 SG311,
penalty= 4

! HEME dummy
CG321  CG2R52 NG2R50 Ni1p       0.0000  0     0.00 ! HEME
CG3C50 CG2R52 NG2R50 Ni1p       0.0000  0     0.00 ! HEME
CG321  CG3C51 NG2R50 Ni1p       0.0000  0     0.00 ! HEME
CG3C51 CG3C51 NG2R50 Ni1p       0.0000  0     0.00 ! HEME
Ni1p   NG2R50 CG3C51 HGA1       0.0000  0     0.00 ! HEME
CG321  CG3C51 NG2R50 Ni1p       0.0000  0     0.00 ! HEME
CG3C51 CG3C51 NG2R50 Ni1p       0.0000  0     0.00 ! HEME
CG3RC1 CG2R52 NG2R50 Ni1p       0.0000  0     0.00 ! HEME
CG2DC1 CG2R52 NG2R50 Ni1p       0.0000  0     0.00 ! HEME
CG2DC1 CG251O NG2D1  Ni1p       0.0000  0     0.00 ! HEME
CG3C51 CG251O NG2D1  Ni1p       0.0000  0     0.00 ! HEME
CG2DC1 CG2R52 NG2R50  Ni1p      0.0000  0     0.00 ! HEME
CG3C51 CG2R52 NG2R50  Ni1p      0.0000  0     0.00 ! HEME
CG3RC1 CG3RC1 NG2R50  Ni1p      0.0000  0     0.00 ! HEME
NG2R53 CG3RC1 NG2R50  Ni1p      0.0000  0     0.00 ! HEME
CG321  CG3RC1 NG2R50  Ni1p      0.0000  0     0.00 ! HEME
CG2R52 NG2R50 Ni1p NG2R50       0.0000  0     0.00 ! HEME
CG2R52 NG2R50 Ni1p NG2D1        0.0000  0     0.00 ! HEME
CG3C51 NG2R50 Ni1p NG2R50       0.0000  0     0.00 ! HEME
CG3C51 NG2R50 Ni1p NG2D1        0.0000  0     0.00 ! HEME
CG251O NG2D1  Ni1p NG2R50       0.0000  0     0.00 ! HEME
CG3RC1 NG2R50 Ni1p NG2R50       0.0000  0     0.00 ! HEME
CG3RC1 NG2R50 Ni1p NG2D1        0.0000  0     0.00 ! HEME

IMPROPERS
! For F430 cofactors
CG2O5  CG2DC1 CG321  OG2D3     88.0000  0     0.00 ! from CG2O5 CG2DC2 CG331 OG2D3, penalty= 0.6
CG2O5  CG2DC1 CG311  OG2D3     88.0000  0     0.00 ! from CG2O5 CG2DC1 CG331 OG2D3, penalty= 1
CG2O3  OG2D2  OG2D2  CG3C51    96.0000  0     0.00 ! from CG2O3 OG2D2 OG2D2 CG321, penalty= 6.5
       CG2DC1 CG2O5  CG2R52 CG251O    96.0000  0     0.00 !
CG251O NG2D1  CG3C51 CG2R53    88.0000  0     0.00 !
CG2R52 NG2R50 CG3C50 CG321     88.0000  0     0.00 !
CG2R52 NG2R50 CG3RC1 CG2DC1    88.0000  0     0.00 !
CG2R52 NG2R50 CG3C51 CG2DC1    88.0000  0     0.00 !
NG2R50 CG2R52 CG3RC1 Ni1p     137.4000  0     0.00 ! HEME
NG2R50 CG3C51 CG2R52 Ni1p     137.4000  0     0.00 ! HEME
NG2D1  CG251O CG251O Ni1p     137.4000  0     0.00 ! HEME
END
RETURN
```

### Supplemental Results

Identification of a modified $F_{430}$ in *Methanosarcina acetivorans*. In previous work,[2] we identified a modified version of $F_{430}$ in the hyperthermophilic *Methanococcales* methanogen, *Methanocaldococcus jannaschii.* Based on the exact mass, characteristic fragment ions, and the UV-vis spectrum, we proposed the modification to be a cyclized mercaptopropionate moiety attached as a thioether to the $17^2$ position of $F_{430}$. To gain insights into the significance and potential function(s) of modified $F_{430}$s, we have explored the existence of modified $F_{430}$s in other methanogens, including a model *Methanosarcinales* methanogen, *Methanosarcina acetivorans*. Interestingly, we identified a modified $F_{430}$ in *M. acetivorans* that is one mass unit less than our previously identified mercaptopropionate-$F_{430}$ (1008 vs. 1009, **Figure S2A**). This difference corresponds to the replacement of a carboxyl group (45 Da) with a primary amide (44 Da). Notably, the mass spectrum of the newly identified $F_{430}$ displays a prominent doubly charged ion (**Figure S2A**), supporting the assignment of an amide-containing modification compared to a carboxyl group. The mass spectrum reveals the characteristic nickel isotope pattern with an intense $[M+2]^+$ peak due to Ni-60 (26% natural abundance) (**Figure S2B**). The modified $F_{430}$ is comprised of four major peaks that elute before the canonical $F_{430}$ during reverse-phase HPLC (**Figure S2C**). The various peaks are expected to be stereoisomers that are likely produced chemically during the processing of the cell extract. UV-vis analysis revealed the characteristic 430 nm absorbance maximum (**Figure S2D**), and the spectrum was identical to that of the unmodified $F_{430}$ in these cells. MS/MS fragmentation shows a major fragment where the modification is cleaved off to yield the parent $F_{430}$ (*m/z* 905), supporting a structure where a single new side-chain to $F_{430}$ has been installed. Based our current spectral data and assuming that this new $F_{430}$ modification is related to our previously reported $F_{430}$ modification,[2] we propose a possible structure with a 3-mercaptopropanamide modification where the sulfur is inserted into a C-H bond. We propose that the modification occurs at the $17^2$ position since this is one of the few sites that would produce the 905 fragment ion and the F-ring appears to be a hotspot for $F_{430}$ modifications.[3,4] However, future detailed structure determination experiments will be required to elucidate the true structure. Interestingly, the modified $F_{430}$ was only identified in *M. acetivorans* cultures grown on acetate and was not present when the organism was grown on methanol or trimethylamine. Although the amount of the modified $F_{430}$ in acetate-grown *M. acetivorans* varies substantially between different preparations, we have observed the modified version existing at amounts up to ~40% of the total $F_{430}$.

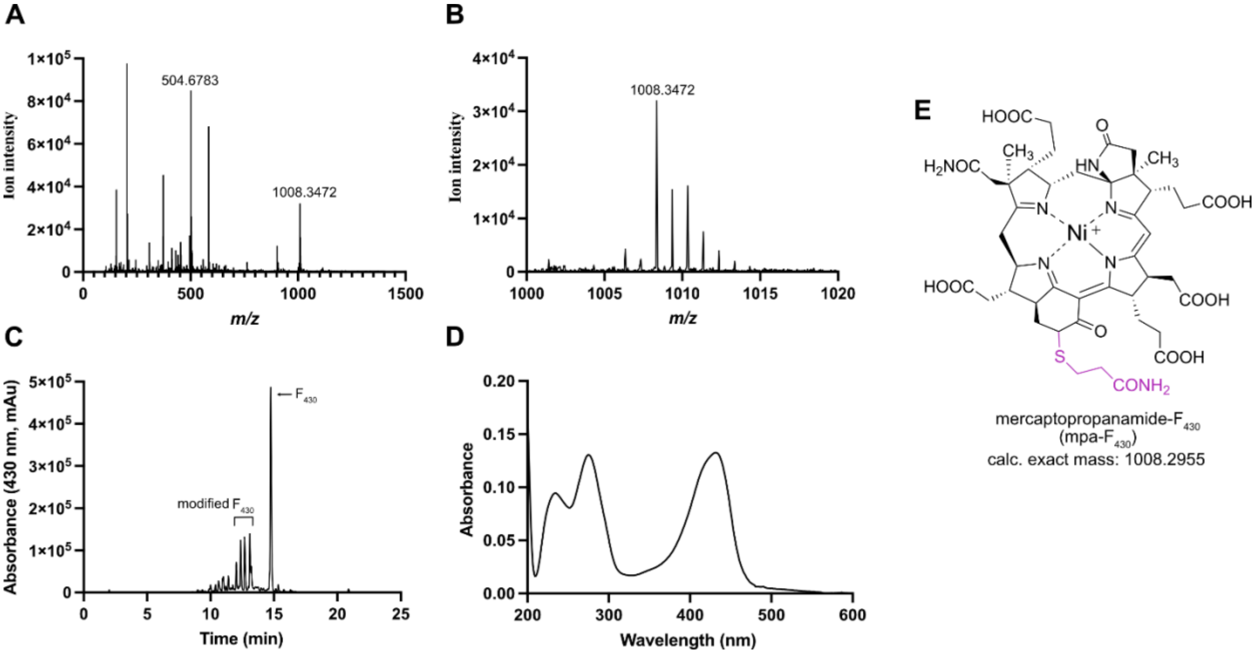

**Figure S2**. **Identification of a modified F430 in *M. acetivorans*.** (A) mass spectrum showing $M^+$ molecular ion at 1008.3472 as well as the prominent doubly charged ion. (B) mass spectrum showing isotope peaks for molecular ion with characteristic nickel isotope peak at $[M+2]^+$. (C) HPLC-DAD analysis with 430 nm extracted chromatogram shown. (D) absorbance spectrum of the modified $F_{430}$. (E) a proposed structure of the modified $F_{430}$.

**Supplemental Tables**

Protonation assignment of MCR residues for MD simulations. The $pK_a$ calculations carried out by the webserver PlayMolecule were visually checked to ensure an optimized hydrogen-bonding network. Crystallographic water molecules and cofactors were kept for the $pK_a$ calculations. The protonation states used in this work are described below for the MCR of *M. acetivorans* (Table S1) and ANME-1 (Table S2).

**Table S1.** Protonation state of His, Glu and Asp residues for *M. acetivorans* MCR in this work.

| Subunit α (chain A/B) | Subunit β (chain C/D) | Subunit γ (chain E/F) |
|---|---|---|
| ASPP353<br>ASPP363<br>ASPP488<br><br>GLUP36<br>GLUP96<br>GLUP291<br><br>HSD73<br>HSE101<br>HSD127<br>HSD145<br>HSE152<br>HSE168<br>HSD223<br>HSD299<br>HSE415<br>HSE454<br>HSE504<br>HSP516 | GLUP 222<br><br>HSD196<br>HSD233<br>HSP362<br>HSE377<br>HSE382 | HSD21<br>HSE43<br>HSD54<br>HSD157<br>HSD159<br>HSE234 |

**Table S2.** Protonation state of His, Glu and Asp residues for ANME-1 MCR in this work.

| Subunit α (chain A/D) | Subunit β (chain B/E) | Subunit γ (chain C/F) |
|---|---|---|
| GLUP396 | GLUP239 | GLUP137 |
| HSD8 | HSD135 | HSE10 |
| HSD84 | HSE152 | HSE42 |
| HSE95 | HSD194 | HSD53 |
| HSD121 | HSE228 | HSD156 |
| HSE146 | HSD232 | HSD158 |
| HSD157 | HSP360 | HSD241 |
| HSE289 | HSE375 | |
| HSE414 | HSE380 | |
| HSD453 | | |
| HSP503 | | |
| HSP519 | | |
| HSD570 | | |
